# Enhanced methionine cycle suppresses naïve CD8^+^ T-cell maturation

**DOI:** 10.1101/2022.08.16.504136

**Authors:** A Saragovi, X Zheng, I Abramovich, L Cohen-Daniel, L Zhuoning, H Raifer, I Omar, A Swisa, M Kuchersky, A Drori, F Marini, O Toker, R Somech, Ch Schmidl, RC Hendrickson, E Gottlieb, M Huber, M Berger

## Abstract

The metabolic pathways controlling naive CD8^+^ T (Tn) cell maturation following thymic egress remain mostly undefined. This is important because immature Tn are a major component of peripheral immune tolerance in newborns and under lymphopenia. In this study we demonstrate that TMRM, a mitochondrial membrane potential marker, could be applied to rapidly identify an immature Tn cell population in the periphery. Applying this marker to perform metabolic and proteomic analysis, we show that immature Tn cells maintain accelerated methionine cycle in respect to mature Tn. This unique metabolic state was associated with restricted Tbx21 locus and diminished immune response in vitro and in vivo. Following our findings, we demonstrate that inhibition of methionine cycle leads to rapid functional maturation of Tn and recovery of immune response to stimuli. Our work provides insight into the way the rate of methionine cycling regulates T cell maturation, opening a path for metabolic manipulation of immune tolerance.

## Introduction

The mechanism governing naive CD8^+^ T-cells (Tn) post-thymic maturation in the periphery is incompletely understood(Boursalian et al., 2004; Cunningham et al., 2018a; Fink, 2013; Hogquist et al., 2015). In particular, the role of metabolism in regulating the transition into mature Tn remains unknown. Understanding the pathways critical for T cell maturation is important because immature Tn are thought to be a major component of immune tolerance adapted to provide protective immunity in newborns and during lymphopenia. (Cunningham et al., 2018a).

Over the last decade, several key studies successfully characterized immature Tn as a distinct subset with unique functional properties. These studies focused on a defined population that recently completed their thymic egress and are therefore highly enriched in immature Tn (Berkley and Fink, 2014a; Cunningham et al., 2018b; Fink and Hendricks, 2011; Friesen et al., 2016a; L. E. Makaroff et al., 2009; Vrisekoop et al., 2008). This immature subset was shown to be generally hyporesponsive to diverse immune stimuli with respect to MN (Boursalian et al., 2004; L. E. Makaroff et al., 2009; Lydia E. Makaroff et al., 2009; Priyadharshini et al., 2010). In contrast, it was established that immature Tn are uniquely adapted to provide protective immEunity under lymphopenic conditions, prevalent for example in neonatal tissue and in lymphodepleted patients (Berkley and Fink, 2014b; Friesen et al., 2016b).

The abundance of immature Tn in the periphery is a function of thymic egress and the rate of maturation. Several factors contributing to Tn maturation and survival have been identified, including IL7R (Kim et al., 2016), the NKAP-associated signaling pathway factor (Hsu et al., 2011), HDAC3 (Hsu et al., 2015), and the zinc finger protein, Zfp335 (Han et al., 2014). Thus, an ensemble of signals and immunological cues regulates Tn maturation.

In activated and effector T cells, metabolic rewiring is a critical regulator of immune function and fate determination (Bantug et al., 2017; Chapman et al., 2020; MacIver et al., 2013; Pearce et al., 2013; Wang and Green, 2012). Changes in mitochondrial mass, morphology, and membrane polarization impact fate determination in activated T cells (Buck et al., 2016; Saragovi et al., 2020; Sukumar et al., 2016a; Vardhana et al., 2020; Yu et al., 2020). Similarly, differences in the levels of glycolysis (Frauwirth et al., 2002; Jacobs et al., 2008), glutaminolysis (Carr et al., 2010; Nakaya et al., 2014) the methionine cycle (Roy et al., 2020; Sinclair et al., 2018), and one carbon metabolism (Ron-Harel et al., 2016) control T cell effector response. Likewise, the abundance of key metabolites including, succinate (Bailis et al., 2020; Nastasi et al., 2021; Tannahill et al., 2013), itaconate (Hooftman et al., 2020), serine (Hooftman et al., 2020) and alanine (Ron-Harel et al., 2019) regulate both inflammation and effector functions. Mature and immature Tn acquire different metabolic signature after activation (Cunningham et al., 2018c, 2017), however, the influence of specific metabolic pathways on the maturation of immature Tn is poorly understood.

Here we describe the use of Tetramethylrhodamine, methyl ester (TMRM)-a mitochondrial membrane polarization stain-as a marker, enabling the identification of two distinct Tn populations: decreased mitochondrial polarization Tn (dpT), and hyperpolarized mitochondria Tn (hpT). Comparative study of the two populations across tissues and mice age cohorts has provided evidence that the hpT are an immature Tn population with increased abundance in the thymus and the periphery of young mice. Additional characterization confirms that hpT differentially respond to immune stimuli and have distinct metabolic network. hpT had increased metabolic drivers of de novo DNA methylation including DNA Methyltransferase 3 (DNMT3), and methionine cycle. ATACseq analysis identified that hpT cells maintain restricted Tbx21 locus in respect to dpT. Likewise, T-bet protein expression was found to be decreased in hpT in relation to dpT. Building on our finding, we demonstrate that inhibition of methionine cycle triggers rapid functional maturation of Tn leading to robust response to immune stimuli both in vitro and in vivo. Our work provides an insight into the way methionine cycleregulates Tn maturation, paving the way for manipulating immune tolerance.

## Results

### Mitochondrial polarization staining distinguishes between two developmental stages of naïve CD8^+^ T cell

Metabolic markers have been used to recognize distinct lymphocyte and monocyte subsets (Katajisto et al., 2015; Romero-Moya et al., 2013; Simsek et al., 2010; Sukumar et al., 2016b). We wanted to characterize a marker that could identify immature naive CD8^+^ T cell populations independent of tissue specificity or time elapsed from thymic output. We chose to use TMRM, a mitochondrial polarization dependent stain, since it is an efficient tool for identification of distinct effector T cell and macrophages populations in humans and mice (Sukumar et al., 2016a). To normalize our findings, we also measured the TMRM intensity of Tn cells following oligomycin (complex V inhibitor) or Carbonyl cyanide-p-trifluoromethoxyphenylhydrazone (FCCP, uncoupler) treatment. These values were used as reference for maximal or minimal TMRM staining, respectively. Flow cytometry analysis of Tn obtained from wild type (WT) mice spleens and stained with TMRM revealed a bipolar TMRM staining pattern, wherein the major subpopulation showed decreased TMRM staining (dpT) and the minor nearly maximal TMRM staining (hpT) (Figuress 1A-B, Figure S1A). To exclude the possibility that the differences observed in polarization state stems from a transient phenomenon, we assessed the capacity of the two Tn subpopulations to regain their original phenotype following mitochondrial membrane uncoupling. Tn cells were treated with FCCP to depolarize their mitochondria and reach minimal TMRM staining. FCCP treated Tn were then washed and allowed to recover from FCCP-induced uncoupling. Following the recovery period, Tn were restrained with TMRM and analyzed by flowcytometry (Figure S1B). We found that after FCCP withdrawal, TMRM staining was identical to that seen in untreated cells (Figures S1C). These results indicate that the differences in TMRM staining pattern observed in Tn is linked to a stable metabolic divergence.

**Figure 1.**
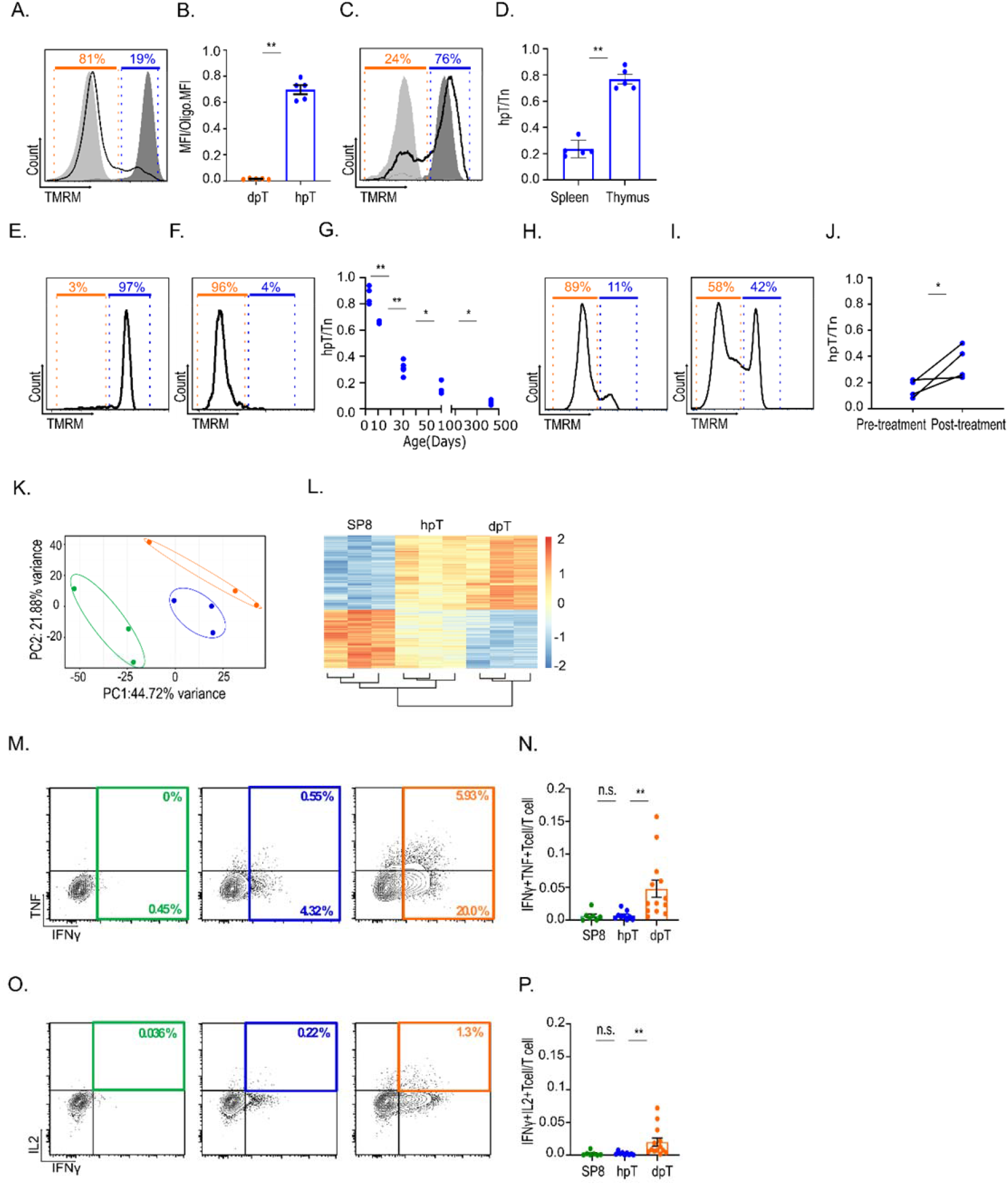
Mitochondrial polarization staining distinguishes between two developmental stages of naïve CD8^+^ T cell: (A-D) Identification of two naïve CD8^+^ T cell (Tn) subsets by TMRM signal quantification, decreased-polarization Tn (dpT – region below orange line) and hyperpolarized Tn (hpT – region below blue line). **A**. Overlay histogram plot of TMRM signal, gated on CD8^+^ CD44^low^ T cells from mouse spleen that were treated with either FCCP (minimal TMRM staining - light gray), oligomycin (maximal TMRM staining - dark gray) or left untreated (black line). **B**. Bar graph, TMRM mean fluorescence intensity (MFI) of dpT or hpT normalized to maximal MFI (oligomycin control group), (n=5 mice). **C**. Same as A., gated on single positive CD8^+^ cells (SP8 - CD8^+^ CD4^-^ CD24^+^ TCRβ^+^) from mouse thymus. **D**. Bar graph, relative abundance of hpT with respect to total Tn or SP8 from mouse spleen or thymus (n=5 mice). **E**. Representative histogram plot, TMRM intensity gated on CD8^+^ CD44^-^ T cells from a spleen of a 1-day old mouse (dpT - region bellow orange line, hpT – region bellow blue line). **F**. Same as in E., from spleen of a 450-day old mouse. **G**. Dot graph, summary of the relative abundance of hpT in mouse spleen from different age cohorts (calculated as the ratio between hpT and Tn), (n=4-5 mice). (H-I) Relative abundance of hpT with respect to total Tn in peripheral blood extracted from 8 week old mice before, or eight days following T cell depletion by I.P. injection with anti-Thy1.2 antibody, dpT (orange) and hpT (blue) **H**. Representative histogram plot, TMRM intensity gated on Tn in peripheral blood of 8 weeks old mice before T cell depletion using I.P. injection of anti-Thy1.2 antibody (dpT - region bellow orange line, hpT -region bellow blue line). **I**. Same as in H., after T cell depletion. **J**. Repeated measure dot graph, summarizing the results in H-I. (n=4 mice). (K-L) ATAC-seq profiling data comparing mouse SP8 thymocytes (green), hpT (blue) and dpT (orange) from spleen. **K**. Principal component analysis. **L**. Heatmap (blue to red represent lower to higher gene accessibility), (N=3 mice). (N-Q) Flow cytometry analysis of intracellular cytokines staining in sorted SP8 thymocytes (green), hpT (blue) and dpT (orange) from spleen, four days after activation using a combination of anti-CD3ε and anti-CD28. **M**. Representative scatter plots of TNF vs. IFN-γ staining. **N**. Bar graph, summary of results presented in N. (n=7-13 mice). **O**. Representative scatter plots of IL-2 vs. IFN-γ staining. **P**. Bar graph, summary of experiment presented in P. (n=7-13 mice). **Statistics** | Mann-Whitney test - P-values - B. ** = 0.0079, D. ** = 0.0079, G. ** = 0.0079, J. * = 0.035 | Kruskal-Wallis test - P-values - N. ** = 0.0051, P. **=0.0077 | error bars represent S.E.M.

To characterize the two Tn populations, we compared them by Forward Scatter (FSC) and Side Scatter (SSC)—proxies for cell size and granularity, respectively—, and saw no differences (Figure S1 D-E). We found that the two Tn populations were indistinguishable in size, or granularity. To examine whether the Tn populations identified are related to differences in T cell receptor (TCR)-repertoire, we analyzed TMRM staining in Tn cells from OT1 or PMEL17, a TCR transgenic mice, giving rise to a single CD8^+^ TCR clone. Similar to WT mice, Tn cells from OT1 and PMEL17 mice gave rise to two distinct subpopulations (Figure S1 F-G).

Naïve T cell populations vary between compartments (Van Den Broek et al., 2018). We therefore next compared the proportion of hpT and dpT cells in secondary and primary lymphatic tissues. We found that -similar to the spleen-Tn from secondary lymph nodes and blood had a majority of dpT cells (Figure 1A-B and S1H-I, S1K). In contrast, the vast majority of thymic single positive CD8^+^ cells (SP8) had hyperpolarized mitochondria (Figure 1C-D, S1J-K).

To examine the relevance of our findings to human immunity, we next analyzed TMRM staining of human T cells. As observed in mice, human resting Tn cells (CD28^hi^, CD45RA^+^) gave rise to bipolar TMRM staining (Figures S1L, S1N), and staining was higher in thymocytes than Tn from peripheral blood or adenoids (Figure S1M-N). Together these findings suggest that as in mice, human hpT relative abundance relates to the rate of thymic output.

Age-related thymic regression is associated with a decline in Tn cell output (George and Ritter, 1996; Lynch et al., 2009; Ma et al., 2013). To support the link between hpT and thymic output we examined the ratio of hpT to total Tn in mice grouped by age. We found that in newborn mice, hpT are the majority (∼80%) of the total Tn cells. This population markedly declined with age, reaching approximately 5% of the Tn cell population in aged mice (Figure 1E-G). In line with these results, the hpT/Tn ratio substantially increased in adult mice following lymphocytes recovery from T cell depletion (Figure 1H-J).

Based on our results, we hypothesized that TMRM staining corresponded to Tn maturation state. We used ATAC-seq to compare the chromatin landscape of SP8 thymocytes, hpT and dpT cells. Both principal component analysis (Figure 1K) and gene clustering analysis (Figure 1L) revealed distinct chromatin accessibility landscapes among the cells, with hpT cells intermediate between SP8 thymocytes and dpT cells.

To assess whether the hpT also shares functional similarities with their thymic progenitors, we evaluated cytokine production of SP8 thymocytes, hpT and dpT in response to activation using anti-CD3ε and anti-CD28 agonist antibodies. In line with the ATAC-seq data, the percentages of cytokine-producing hpT and SP8 thymocytes were similar and reduced in comparison to cytokine-producing dpT (Figure 1M-P). Overall, our results so far indicate that TMRM-based mitochondrial polarization staining distinguishes between immature-hpT and mature-dpT naïve CD8^+^ T cell subsets.

### Immature-hpT are hyporesponsive to diverse stimuli relative to mature-dpT

Immature CD8^+^ T cells support protective immunity in lymphopenic environment and during chronic inflammations due to their reduced responsiveness to diverse immune stimuli (Berkley and Fink, 2014b; Cunningham et al., 2018a; Friesen et al., 2016b). We therefore thought to compare the functional properties of the immature-hpT in respect to the mature-dpT both *in vitro* and *in vivo*.

To ensure that T cell functional assays are not influenced by TMRM related toxicity, we measured Tn activation *in vitro* in the presence or absence of high TMRM concentration (100 μM). In line with previous reports, the presence of TMRM in the medium had no effect on cell proliferation or elevation of activation markers (Figures S2A-B). Next, we compared the response of both Tn subsets to homeostatic proliferation signals. Immature-hpT and mature-dpT cells isolated from congenic Ly5.2 and Ly5.1 mice, respectively, were adoptively transferred at 1:1 ratio into *Rag1*^*-/-*^ recipient mice and monitored for proliferation in the peripheral blood at different time points (Figure 2A). Five days following the transfer, dpT-donor cells (Ly5.1) rapidly increased their abundance in the total Tn-donor pool relative to hpT-donor cells (Ly5.2) (Figure 2B). This ratio between the two competing donor subsets remained constant also twenty-five days after the transfer (Figure 2B). These results suggest that the mature-dpT responds better to homeostatic expansion cues relative to the immature-hpT.

**Figure 2.**
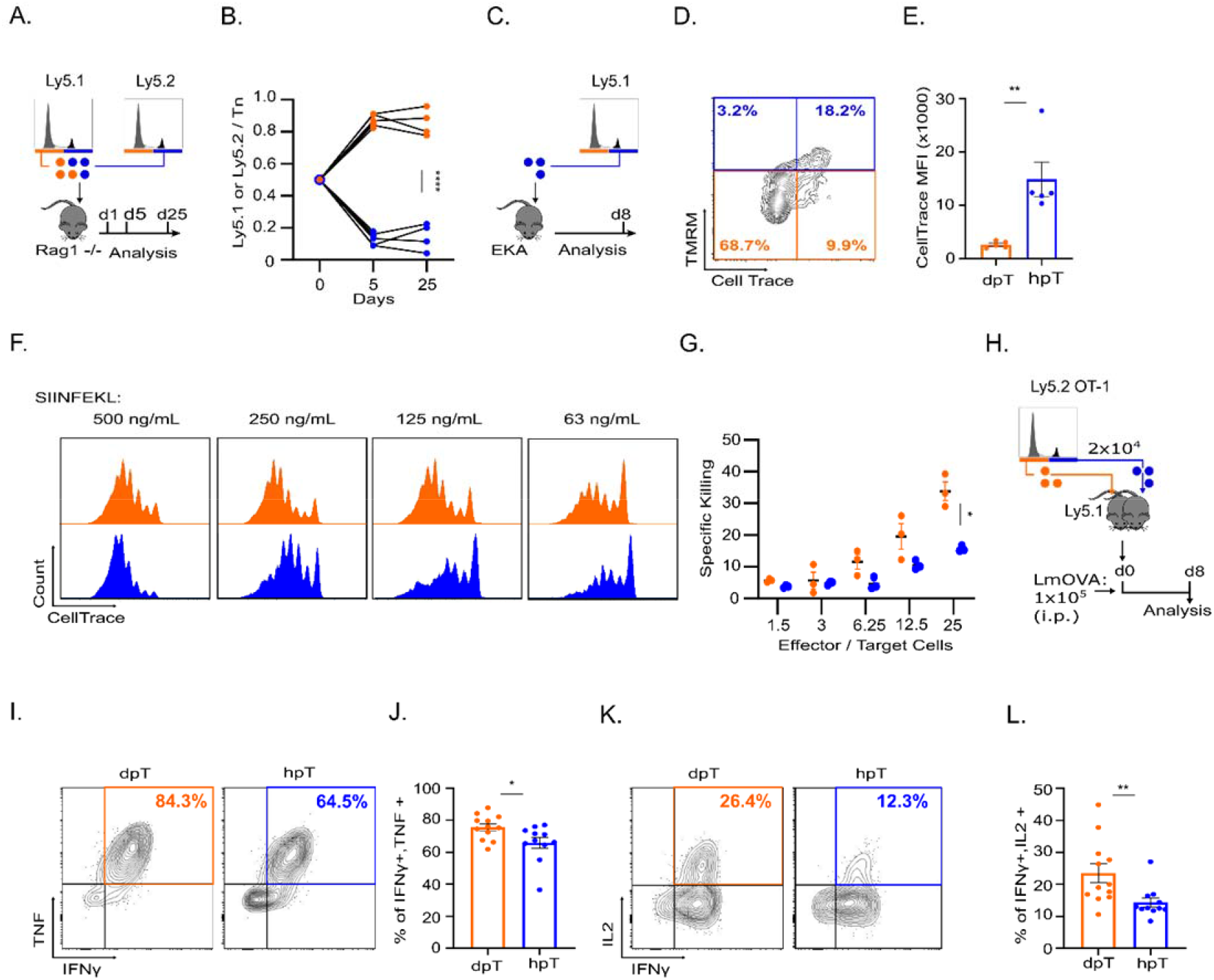
Immature-hpT are hyporesponsive to diverse stimuli relative to mature-dpT both *in vitro* and *in vivo*: (A-B) hpT and dpT cells isolated from syngeneic Ly5.2 and Ly5.1 mice, respectively, were adoptively transferred at 1:1 ratio into *Rag1*^*–/–*^ recipient mice and monitored in the blood by flow cytometry at the indicated time points. **A**. Experiment schematics. **B**. Repeated measure dot graph, the percentage of each subset from the total Tn population (blue dots: hpT-originated cells from Ly5.2 mice; orange: dpT-originated cells from Ly5.1 mice). (n=4 mice). (C-E) Cell Trace labeled isolated hpT from OT1/Ly5.1 transgenic mice were adoptively transferred into semi-lymphopenic, elektra/Ly5.2 mice. Eight days after the adoptive transfer splenocytes from the recipient mice were stained with TMRM, anti-CD8 and anti-CD45.1 antibodies and subjected to flow cytometer analysis (n=4 mice). **C**. Experiment schematics. **D**. Representative scatter plot of TMRM and Cell Trace staining gated on CD8^+^ CD45.1^+^ T cells. **E**. Bar graph, Cell Trace intensity of the TMRM^high^ (blue gate) and TMRM^low^ (orange gate) populations shown in D (n=5 mice). **F**. Representative overlay histogram, plot of Cell Trace intensity of dpT (orange) and dpT (blue) isolated from OT1/Ly5.1 spleen and activated for 72 hours with SIINFEKL at the indicated concentrations (n=3 biological replicates per concentration). **G**. Scatter plot, *ex vivo* killing assay comparing isolated dpT derived effector cells (orange dots) and hpT derived effector cells (blue dots). Isolated dpT and hpT that were activated with anti-CD3ε (0.05 μg/well) and anti-CD28 (0.05 μg/well) for 5 hours were incubated with [S35]-Methionine labeled target cells (P815) at the indicated ratios for 4 hours. Radioactivity levels of the medium were measured by beta-counter. Specific killing was calculated as follows: ((radioactive reading – spontaneous release) / (total l35abeling – spontaneous release))*100, (n=3 biological replicates per ratio group) (H-K) (2×10^4^ hpT or dpT cells isolated from syngeneic OT-1 Ly5.2 mice were adoptively transferred into LmOVA (10^5^ cfu) infected Ly5.1 recipients. Eight days post transfer CD8^+^CD45.2^+^ cells were analyzed for cytokines secretion by intracellular staining. **H**. Experiment schematics. **I**. Representative scatter plots showing TNFα vs. IFNγ intensities in dpT and hpT **J**. Summary of results presented in I. (n=12 mice) **K**. Representative scatter plots showing IL2 vs. IFNγ intensities in dpT and hpT **L**. Summary of results presented in K (n=12 mice). **Statistics** | Mixed effect analysis - P-values - B. **** < 0.0001, G. *= 0.0258 | Mann-Whitney test - P-values -, E. **=0.0079, J. *=0.0215, L. **=0.0048 | error bars represent s.e.m..

To further explore the lag in the immature-hpT response in a lymphopenic environment, we followed the proliferation and mitochondrial membrane polarization levels of hpT-donor cells in a mildly lymphopenic environment *in vivo*. Isolated Cell-Trace labeled hpT cells were adoptively transferred to semi-lymphopenic, *elektra* (Berger et al., 2010) mice for 8 days. Donor cells were then analyzed by flow cytometry to correlate the levels of proliferation and TMRM intensity (Figure 2C). In line with our hypothesis, hpT-donor cells that depolarized their mitochondria, proliferated to a greater extent than hpT-donor cells that retained their increased mitochondrial polarization levels (Figure 2D-E).

Next, we tested whether the immature-hpT population also maintains a reduced responsiveness to activation stimuli compared to that of the mature-dpT population. Isolated Cell-Trace labeled hpT and dpT cells from the spleens of OT1 transgenic mice were stimulated with various concentrations of ovalbumin-derived SIINFEKL peptide. We found that mature-dpT cells proliferate at a significantly higher rate in a peptide-concentration-dependent manner in comparison to the immature-hpT (Figure 2F). Similarly, the killing capacity of the dpT population was substantially higher than that of its hpT counterpart, reaching a two-fold increase at most ratios of effector *vs*. target cells (Figure 2G).

To substantiate our findings *in vivo*, we examined the potential of the two Tn subsets to acquire effector functions upon infection with an intracellular bacterium *Listeria monocytogenes* (Lm) strain recombinant for chicken ovalbumin (LmOVA), (Figure 2H). We conducted transfers of OVA-specific hpT OT1 cells (CD45.2) or dpT OT1 cells (CD45.2), which were injected into congenic WT mice (CD45.1). To activate the donor cells *in vivo*, the recipient WT mice were infected with LmOVA. The capabilities of the transferred cells to acquire effector functions were examined eight days after infection by intra cellular staining for IFN-γ, TNF and IL-2 (Figure 2K). In line with our *in vitro* findings, higher percentages of IFN-γ^+^TNF^+^ double positive (Figure 2I-J, S3A), and IFN-γ^+^IL-2^+^ double positive (Figure 2K-L) were present in dpT-donor cells in comparison to hpT cells. Similar to previously reported immature Tn subsets (Boursalian et al., 2004; L. E. Makaroff et al., 2009; Lydia E. Makaroff et al., 2009; Priyadharshini et al., 2010), our findings suggest that the immature-hpT population entails delayed response both to activating and to lymphopenic stimuli, and a lower potential to differentiate into “canonical” cytotoxic T lymphocytes (CTLs) fate.

### Immature-hpT maintain distinct metabolic profile in comparison to mature-dpT

Differences in mitochondrial membrane potential are linked to variation in mitochondrial morphology and metabolic networks in both monocytes and lymphocytes (Buck et al., 2016; Sukumar et al., 2016a; Yu et al., 2020). To compare the mitochondrial profile of the immature-hpT population to the one of the mature-dpT, we used the mtDendra2 transgenic mice system, which enables the evaluation of difference in both mitochondrial mass and morphology (Pham et al., 2012; Ron-Harel et al., 2016; Saragovi et al., 2020). Comparison of mtDendra2 in the two Tn populations revealed decreased mitochondrial mass in the immature-hpT relative to the mature-dpT (Figure 3A). Similarly, ImageStream based analysis of the mitochondria showed a different mitochondrial distribution between the two Tn populations (Figure S4A-C). In line with these results, Seahorse-oxygen dependent fluorescence-based assessment of mitochondrial respiration-in both populations demonstrated that the immature-hpT maintain lower Spare Respiratory Capacity (SRC) in respect to the mature-dpT (Figure 3B). Finally, we used proteomic analysis to quantify the levels of mitochondrial-associated proteins, in both immature-hpT and mature-dpT. We identified seventeen enriched mitochondrial associated proteins in dpT vs. only four in hpT, demonstrating that mature-hpT maintain increased levels of mitochondrial associated proteins in respect to the immature-hpT population (Figure 3C). Together these observations demonstrate that the immature-hpT hold decreased mitochondrial mass and respiratory capacity in respect to the mature-dpT.

**Figure 3.**
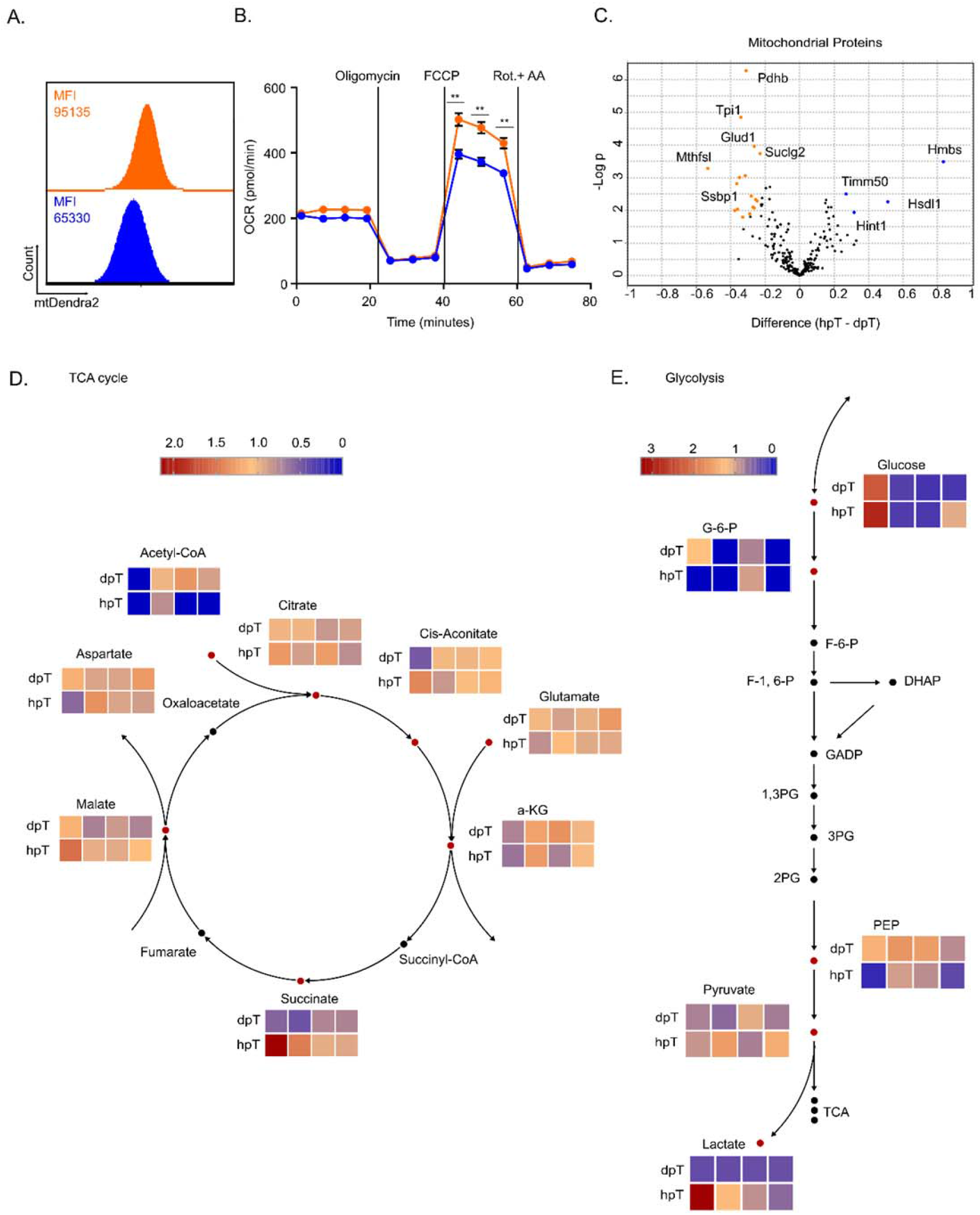
hpT maintain distinct metabolic profile in respect to dpT: **A**. Representative overlay of Dendra2 intensity, gated on CD8^+^ CD44^low^ extracted from spleens of mtDendra2 transgenic mice spleen, dpT (orange) and hpT (blue). **B**. Continuous dot graph, oxygen consumption rate (OCR) measured by seahorse XF24, following consecutive injections of Oligomycin [1µM], FCCP [20µM], Rotenone + Antimycin-A [1 µM] (Rot+AA) of isolateed dpT (orange) and hpT (blue) at the indicated time points (n=7 biological replicates) **C**. Volcano plot of mitochondria-associated proteins of hpT and vs. dpT isolated from mouse spleens. Orange and blue dots mark ANOVA significant proteins that were significantly elevated in dpT and hpT respectively (n=5 biological replicates from a pool of 10 mice). **D**. TCA cycle related metabolic profile of hpT and dpT (n=4 mice). **E**. Glycolysis related metabolic profile of hpT and dpT (n=4 mice). **Statistics** | Mann-Whitney test - P-values -, B. **= 0.0012. | ANOVA test - P-values – C. at the indicated p values | error bars represent s.e.m

Next, we examined the metabolic profile of each population using metabolomic analysis. We focused on metabolic circuits that were previously shown to be important to T cell function and fate, including TCA and glycolysis. We found higher abundance of Acetyl-CoA in the mature-dpT in respect to the immature-hpT, a metabolite associated with increased mitochondrial activity and acetylation events (Pietrocola et al., 2015). In contrast, succinate, a metabolite which was previously reported to stabilize HIF1a (Tannahill et al., 2013), was higher in the immature-hpT in comparison to the mature-dpT (Figure 3D). Analysis of the glycolytic pathway, a hallmark of T cell biology (Fox et al., 2005; Gerriets and Rathmell, 2012), revealed differences in two intermediates between the two populations. Specifically, we found that phosphoenolpyruvate, a metabolite important for effector T cell function (Ho et al., 2015), was higher in the mature-dpT populations (Figure 3E). In contrast, both cellular and secreted lactate were higher in the immature-hpT population (Figure 3E, S4D). Collectively, these findings demonstrate that the immature-hpT maintains a distinct metabolic network in respect to the mature-dpT subset.

### Methionine cycle and Dnmt activity are increased in immature-hpT in respect to mature-dpT

We next attempted to identify a link between the functional and the metabolic differences observed between the two Tn subsets. Further analysis of proteomics data from the two Tn subsets highlighted several key metabolic circuits that are known to affect cellular function (Figure 4A-B). As expected, mitochondrial, electron transport chain and fatty acid oxidation proteins were enriched in the mature-dpT population in respect to the immature-hpT. Likewise, proteins related to the amino acid metabolism and the folate cycle, shown to be critical for T cell activation (Miyajima, 2020), were also enriched in the mature-dpT subset (Figure 4A-B). In contrast, the immature-hpT demonstrated higher enrichment of proteins related to glycolysis, nucleotide metabolism and the methionine cycle (Figure 4A-B).

**Figure 4.**
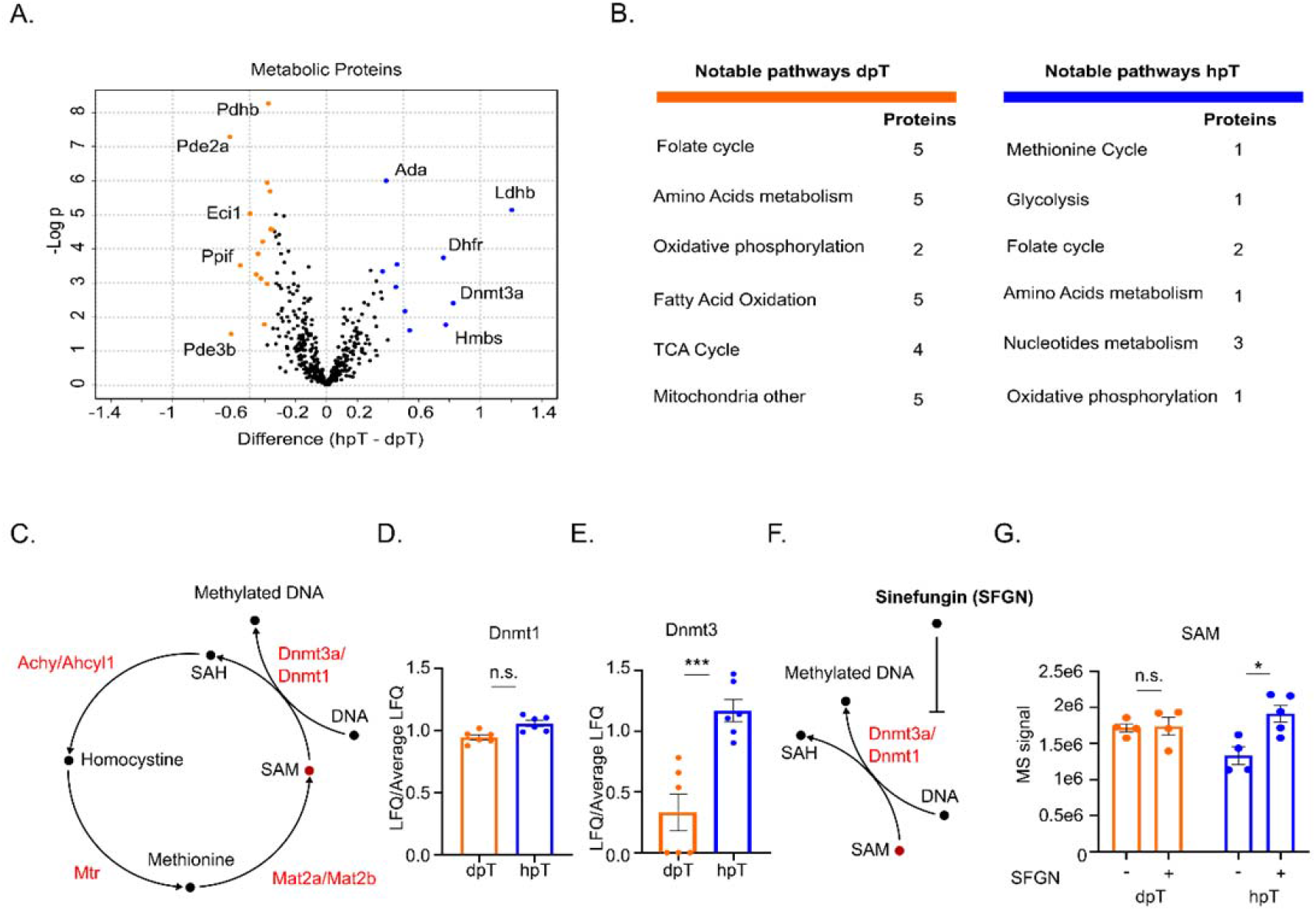
Methionine cycle and Dnmt3a activity are accelerated in immature-hpT in respect to mature-dpT: **A**. Volcano plot of proteins associated with metabolic pathways detected by proteomic analysis of hpT vs. dpT isolated from mouse spleens. Orange and blue dots mark significant proteins that were elevated in dpT and hpT, respectively (n=5 biological replicates from a pool of 10 mice). **B**. Summary of A., notable metabolic pathways presented with substantial differences in protein expression between dpT and hpT. **C**. Schematic of the methionine cycle **D**. Relative protein expression level of Dnmt1 in hpT (orange) and dpT (blue) (n=5-6 biological replicates from a pool of 10 mice). **E**. Relative protein expression level of Dnmt3 in hpT (orange) and dpT (blue) (n=5-6 biological replicates from a pool of 10 mice). **F**. Schematic of Sinefungin inhibition of SAM-dependent methyltransferases. **G**. metabolomics analysis of SAM levels in dpT (orange) and hpT (blue) that left untreated or treated with Sinefungin for 12 hours. **Statistics** | ANOVA test - P-values – A. at the indicated p values | Mann-Whitney test - P-values – E. n.s., ***=0.0001, G. *= 0.0317| error bars represent s.e.m

The methionine cycle plays a central role in determining cell fate (Shiraki et al., 2014). The conversion of S-Adenosyl-L-methionine (SAM) to S-Adenosyl-L-homocysteine (SAH) in the methionine cycle by DNA methyltransferases and specifically DNMT3a is coupled with *de novo* DNA methylation. Thus, the rate by which SAM is converted to SAH is a dominant determinant of cellular DNA methylation (Goll and Bestor, 2005; Zhang, 2018) (Figure 4C).

Given the increased levels of DNMT3a protein in the immature-hpT population (Figure 4D-E), we next examined whether the immature population had increased DNMT activity. To directly measure the rate of SAM to SAH conversion we compared the levels of methionine cycle intermediates extracted from the two Tn populations. As the methionine cycle is linked to several pathways, we determined the rate of SAM to SAH by analyzing the changes in SAM levels following introduction of Sinefungin, a SAM analog, which inhibits methyltransferases activity (Gros et al., 2013; Shao et al., 2017). Metabolomic analysis of SAM prior to treatment has shown an increased level of the intermediate in the mature-dpT cells in respect to immature-hpT. However, following Sinefungin treatment we observed a significant increase in the level of SAM in the immature-hpT but not in the mature-dpT (Figure 4F-G). The increase in the levels of SAM following the inhibition of DNMTs demonstrates that under normal conditions the immature-hpT maintain higher levels of DNMT activity.

### Reduced chromatin accessibility at Tbx21 locus associates with decreased T-BET expression in immature-hpT cells

Differences in methionine cycle and DNMTs activity may be linked to variation in chromatin modification and gene accessibility. We therefore searched for differences in the chromatin accessibility of key functional clusters in the two Tn subsets. Because our analysis also pointed at other possible regulators of gene expression, we chose to use ATAC-seq to also capture other possible effects of chromatin regulators observed including histone acetylation and methylation.

ATAC-seq analysis of closed chromatin regions in sorted mature-dpT have identified two gene clusters with reduced chromatin accessibility. The largest cluster restricted in dpT, included 31 genes (Figure 5A-B) linked to EGFR including Vav2, Vav3, Cd24a, Ccr4 and other inhibitory or developmental genes (Figure S5A). Notably a second smaller cluster included a number of mitochondrial genes (Figure 5A-B S5A). The reduced accessibility of mitochondrial genes in mature-dpT in respect to immature-hpT may suggest that mitochondrial biogenesis is linked to Tn maturation process. ATAC-seq analysis of the immature-hpT population has revealed only one restricted gene cluster. This gene cluster included several genes central for T cell activation and cytokine secretion. These included the transcription factor Tbx21 and the cytokine regulator Socs3 (Figure 5C-D, S5A) as well as Trib1 and Cebpb, which are critical for the regulation of T cell function (Berberich-Siebelt et al., 2000; Rome et al., 2020).

**Figure 5.**
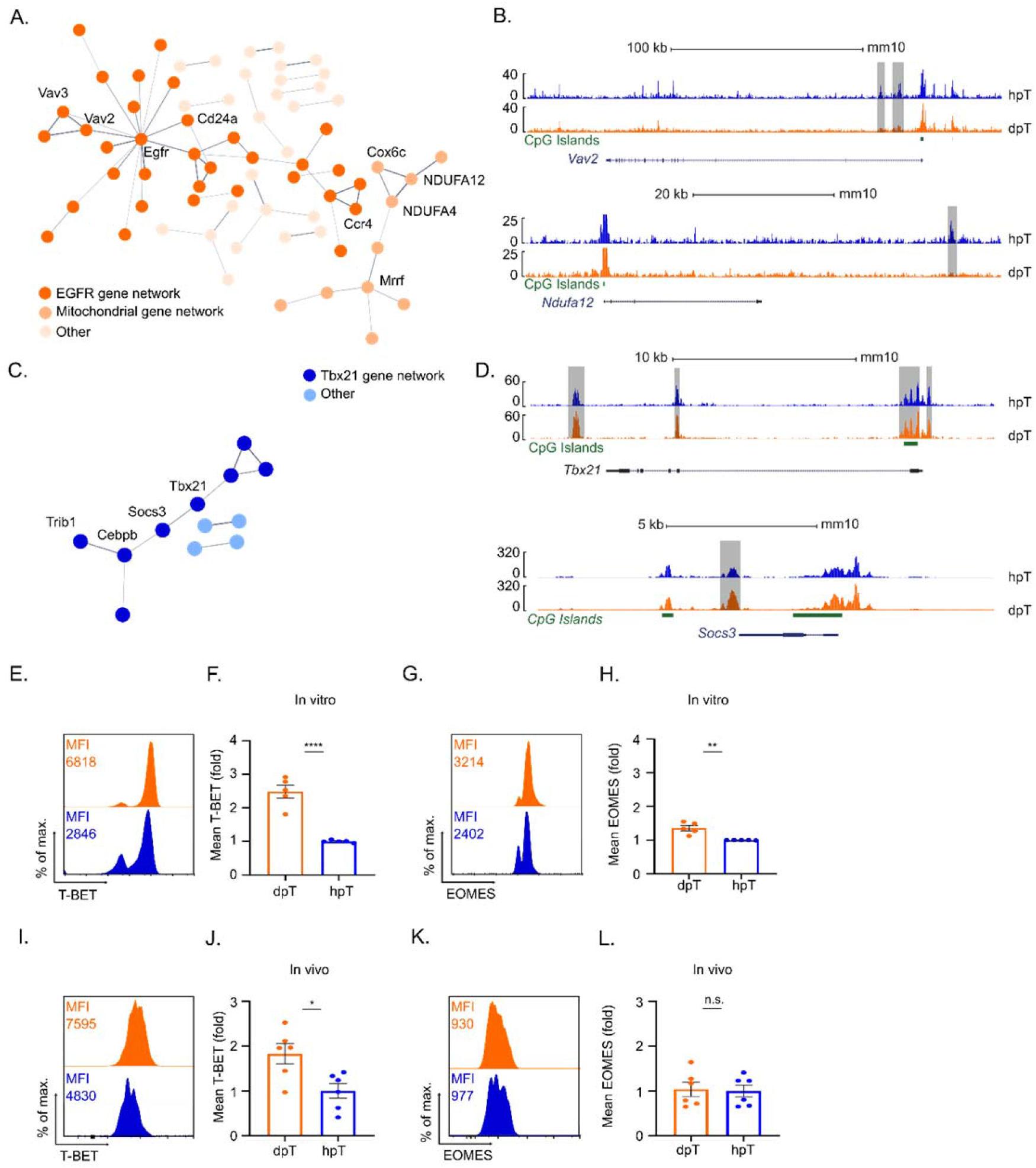
Reduced chromatin accessibility at Tbx21 locus correlates with decreased T-BET expression in immature-hpT cells: (A-D) ATAC-seq analysis of hpT and dpT **A**. String based genes network analysis of restricted gene locus in dpT (n=3 mice). **B**. Sample tracks from each of the restricted networks presented in A. **C**. String based genes network analysis of restricted gene locus in hpT. (n=3 mice). **D**. Sample tracks from each of the restricted networks presented in C. (E-H) dpT and hpT isolated from mice spleens were activated *in vitro*. Ninety-six hours post activation cells were permeabilized and analyzed by flow cytometry for different transcription factors. **E**. Representative overlay histogram of T-bet expression in dpT (orange) and hpT (blue). **F**. Bar graph, summary of results presented in E (n=5 mice). **G**. Representative overlay histogram of EOMES expression in dpT (orange) and hpT (blue). **H**. Bar graph, summary of experiment presented in E (n=5 mice). (I-L) dpT and hpT isolated from Ly5.2 OT1 mice were adoptively transferred into Lm-OVA infected Ly5.1 mice. Eight days post transfer, CD45.2^+^ CD8^+^ cells were permeabilized and analyzed by flow cytometry for the expression levels of T-BET and EOMES. **I**. Representative overlay histogram of T-bet expression in dpT (orange) and hpT (blue). **J**. Bar graph, summary of results presented in I (n=6 mice). **K**. Representative overlay histogram of EOMES expression in dpT (orange) and hpT (blue). **L**. Bar graph, summary of results presented in K (n=6 mice). **Statistics** | Mann-Whitney test - P-values – F. ****<0.0001, H. **=0.0016, J. *=0.026, L. n.s. | error bars represent s.e.m

To verify our ATAC-seq findings, we measured the levels of both T-BET (encoded by Tbx21), and the T cell regulator EOMES, classified under the same family of transcription factors in both immature-hpT and mature-dpT following activation with CD3/CD28 *in vitro*. We found that during T cell differentiation *in vitro* T-BET protein expression levels were over two-fold higher in mature-dpT in respect to immature-hpT (Figure 5E-F). In a similar manner, but to a lesser extent, we observed significantly higher levels of EOMES in dpT in respect to hpT (Figure 5G-H). To confirm our finding in a more physiologically relevant system, we measured the protein expression levels of both T-BET and EOMES following LmOVA infection. Sorted hpT and dpT from OT1 mice were adoptively transferred to congenic WT mice, which were then infected with LmOVA. Eight days following LmOVA infection congenic effector cells derived from either immature-hpT or mature-dpT were analyzed by flow cytometry. As we observed *in vitro*, LmOVA induced effector hpT expressed significantly lower levels of T-BET in respect to effector dpT (Figure 5I-J). We found no significant differences in EOMES expression *in vivo* suggesting that EOMES differences might be short-lived (Figure 5K-L). Together these findings point at a possible link between the unique metabolic network of each of the subsets and the accessibility of loci critical for CD8^+^ T cell terminal differentiation into effector cells (Duckworth et al., 2021; Joshi et al., 2007; Kaech and Cui, 2012).

### Inhibition of methionine cycle and DNA methyltransferase elevates T-BET expression and activation propensity of immature-hpT

Tbx21 expression level was previously linked to DNMT3a activity in T cells (Abdelsamed et al., 2020; Herek et al., 2020; Ladle et al., 2016; Pham et al., 2013; Q. Yu et al., 2012). We therefore thought to explore a possible causal link between the increased methionine cycle activity observed in immature-hpT and the reduced *Tbx21* accessibility and T-BET expression. To this end, we activated immature-hpT with or without an effective dose of Sinefungin (SFGN). Providing that Tbx21 expression is suppressed by DNMT activity, we hypothesized that the inhibition of DNMT and methionine cycle will lead to increased expression of T-BET in immature-hpT. Flow cytometry analysis of T-BET expression in activated hpT revealed increased T-BET protein expression in SFGN treated in respect to untreated cells *in vitro* (Figure 6A-B). In contrast, SFGN treatment did not affect the expression levels of EOMES in activated hpT (Figure 6C-D).

**Figure 6.**
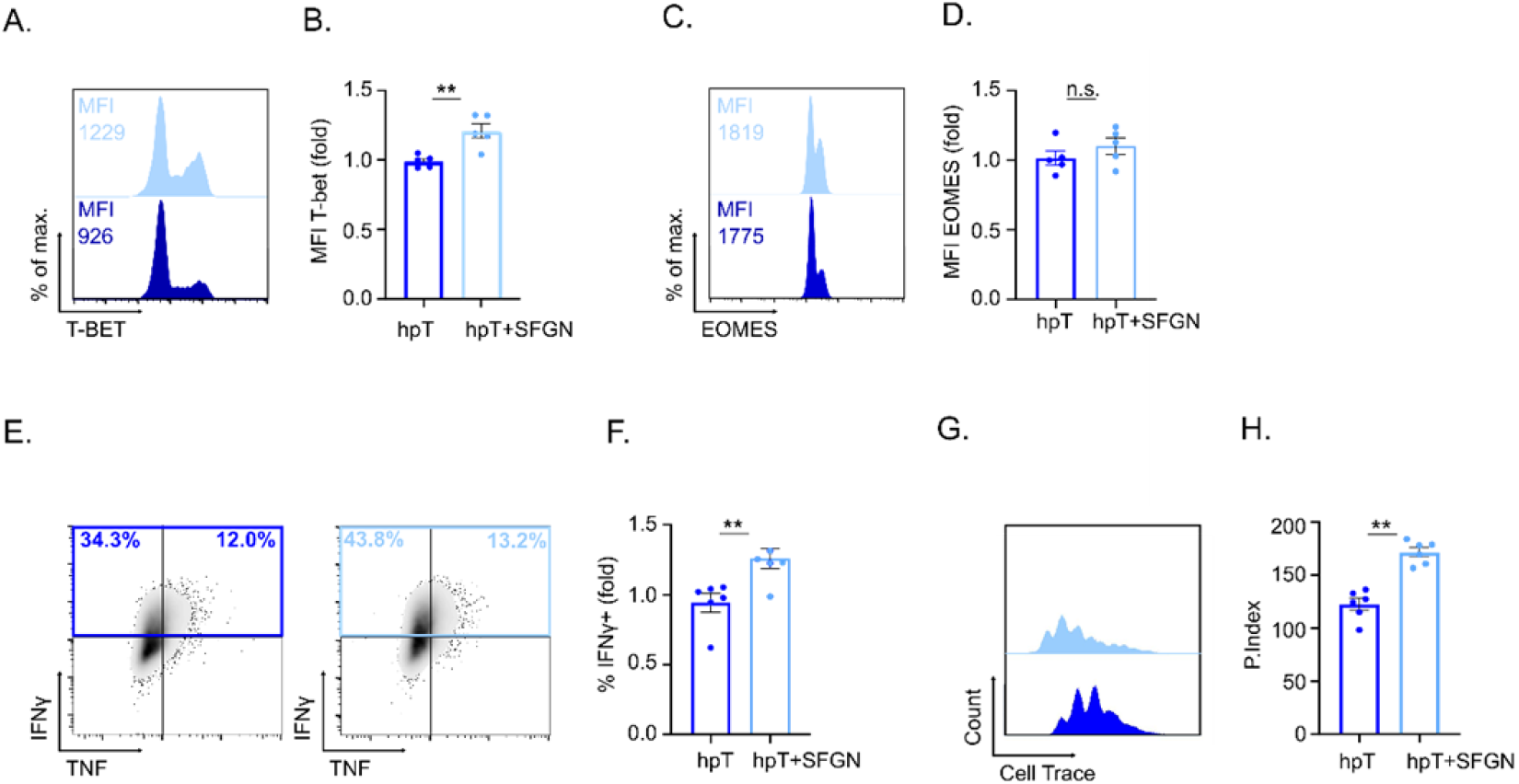
Inhibition of methionine cycle and DNA methyltransferase elevates T-Bet expression and activation propensity of immature-hpT: (A-D) Isolated hpT from mice spleens were activated *in vitro* with (light blue) or without (blue) 30 µM Sinefungin (SFGN). Ninety hours post activation cells were permeabilized and analyzed by flow cytometry for the expression levels of T-BET and EOMES. **A**. Representative overlay histogram of T-BET expression in hpT (blue) or hpT treated with Sinefungin (light blue). **B**. Bar graph, summary of results presented in A (n=5 mice). **C**. Representative overlay histogram of EOMES in hpT (blue) or hpT treated with Sinefungin (light blue). **D**. Bar graph, summary of results presented in C (n=5 mice). (E-F) Isolated hpT from mice spleens were activated *in vitro* with (light blue) or without (blue) 30uM Sinefungin. Ninety-six hours post activation cells were permeabilized and analyzed by flow cytometry for expression levels of IFNγ and TNFα. **E**. Representative scatter plot of TNFα and IFNγ in hpT (left panel) or hpT treated with SFGN (right panel). **F**. Bar graph, summary of results presented in E (n=5-6 mice). (G-H) Cell trace labeled Tn were activated *in vitro* without (blue) or with (light blue) 30 µM Sinefungin. Seventy-two hours post activation CD8^+^ cells were analyzed by flow cytometry for Cell Trace intensity **G**. Representative overlay histogram of Cell Trace signal in control Tn (blue) or SFGN treated Tn (light blue). **H**. Bar graph, summary of experiment presented in G. Proliferation Index (P. Index) calculated as sum (generation * % cell frequency), (n=6 mice). **Statistics** | Mann-Whitney test - P-values – B. **=0.0079, D. n.s., F. **=0.0096, H. **=0.0022. | error bars represent s.e.m

To test whether SFGN based inhibition of DNMT also affects immature-hpT function, we examined cytokines secretion in hpT stimulated in the presence of SFGN or left untreated. Analysis of INF-γ and TNF in hpT following anti-CD3/CD28 stimulation identified a significant increase in SFGN treated hpT in respect to the untreated control (Figure 6E-F). Finally, to evaluate the effect of SFGN as a tool to modulate Tn proliferative capacity, we activated CellTrace labeled mouse Tn in the presence or absence of SFGN. In line with our previous results, SFGN treatment led to a substantial increase in the proliferation of activated CD8^+^ T cells (Figure 6G-H), providing further evidence for a causal link between methionine cycle-DNMT3 activity and the reduced responsiveness observed in the immature Tn cells. Together our findings demonstrate that accelerated methionine cycle and DNMT activity maintain naive CD8^+^ T cell hyporesponsiveness during their maturation process.

## Discussion

Immature Tn have improved capacity to provide protection during chronic viral inflammation (Berkley and Fink, 2014b; Friesen et al., 2016b). Despite their pivotal part in chronic viral inflammation, very little is known about the factors and processes that regulate immature Tn maintenance and maturation. Recently it was suggested that, similar to other immune cells, immature Tn maintain a distinct metabolic network that regulates their maturation (Cunningham et al., 2018c, 2017). However, whether and how immature Tn metabolism regulates their maturation was unknown.

A substantial constraint to investigating Tn maturation process is the luck of simple and available markers, independent of tissue localization or time elapsed from thymic egress. We therefore first investigated whether we could apply TMRM to distinguish immature Tn from the total peripheral Tn population. In mature mice, we identified two populations; the minor subpopulation (hpT), maintained a substantially higher mitochondrial membrane potential relative to the major naïve CD8^+^ T cell subpopulation, dpT. Using numerous assays, we identified a strong positive association between thymic output and the proportions of hpT in the total naïve CD8^+^ T cells pool. We also found that the larger dpT maturates from the hpT progenitor subset. To verify that TMRM could be used as a probe for immature hpT we conducted comparative ATAC-seq profiling of thymic SP8, hpT and dpT splenocytes. This assay revealed that hpTs represents an immature subset in transition between SP8 and mature-dpT. The ability to identify immature-Tn using a simple and commercially available mitochondrial probe, opened an opportunity for us and others to better characterize immature-hpT and establish the cues regulating its abundance and function.

We next applied our system to examine the functional differences between immature-hpT and mature-dpT. Comparative functional analysis demonstrated that the immature-hpT subset has a lower propensity to respond to various immune stimuli *in vitro* in respect to mature-dpT. Similarly, hpT showed reduced ability to acquire effector functions and secret cytokines following activation *in vivo*. These findings are aligned with previous literature, which demonstrated that a key property of immature-Tn is general hyporesponsiveness to diverse immune stimuli (Boursalian et al., 2004; L. E. Makaroff et al., 2009; Lydia E. Makaroff et al., 2009; Priyadharshini et al., 2010).

To inspect whether hpT functional hyporesponsiveness is related to distinct metabolic networks, we quantified both metabolic and mitochondria related proteins as well as metabolites extracted from immature-hpT and mature-dpT. Likewise, we tested mitochondrial capacity using mtDendra2 mice, quantification of mtProteins and Seahorse analysis. We found that immature-hpT maintain decreased mitochondrial mass, proteins, and respiratory capacity in respect to mature-dpT. Proteomics and metabolomics analysis has highlighted a marked difference in the metabolic network in the two subsets. Immature-hpT had increased levels of succinate, a proinflammatory metabolite, and lactate in comparison to mature-dpT. These findings suggest that mitochondrial biogenesis and rewiring, which are critical for T cell activation, may also be important for Tn maturation.

The mature-dpT subset maintained increased levels of PEP, a metabolite associated with improved effector function (Ho et al., 2015). In addition, we found an increased acetyl-CoA abundance in mature-dpTs relative to immature-hpTs. Acetyl-CoA is central for chromatin and protein acetylation and may be central for acquiring the mature phenotype. In addition, our analysis revealed an increase in abundance of DNMT3, a methionine cycle enzyme, in immature-hpT relative to mature-dpT. Motivated by this finding, we examined the rate of methionine cycle by SFGN inhibition. Following inhibition, we observed a significant increase in SAM signal in treated immature-hpT in comparison to untreated immature-hpT control. Under similar conditions, mature-dpT demonstrated no significant change in SAM signal in respect to control.

Acetyl-CoA and SAM-DNMT are both involved in chromatin modulation (Goll and Bestor, 2005; Pietrocola et al., 2015). ATAC-seq analysis of chromatin restriction sites in the two subsets had revealed several differences between the two populations. Mature-dpT had two major restricted gene networks; The Vav-EGFR constellation, an inhibitory signaling pathway, a well-established regulator of immune activity; and the mitochondrial constellation. The lower accessibility of mitochondrial genes in the mature-dpT may imply that these cells have reached the mitochondrial mass required for normal mature Tn function.

In immature-hpT, we identified one restricted gene constellation consisting of Tbx21 and related genes. The regulation of *Tbx21* was previously linked to DNMT activity (Abdelsamed et al., 2020; Herek et al., 2020; Ladle et al., 2016; Pham et al., 2013; Q. Yu et al., 2012). In line with these reports, we observed a marked increase in T-bet expression in immature-hpT treated with SFGN following activation both *in vitro* and *in vivo*. Consistently, SFGN treatment significantly increased the capacity of immature-hpT to acquire effector function following stimuli.

Overall, our study provides a detailed mechanistic insight into the way methionine cycle rate regulates Tn maturation. As the methionine cycle can be pharmacologically manipulated, our findings open a path toward regulating CD8^+^ T cell tolerance and propensity to response..

## Materials and methods

### Human study approval

Human blood samples were obtained via Shaare Zedek Medical Center Jerusalem, Helsinki committee approval number: 143/14. Human thymus samples were obtained via Sheba Medical Center, Tel Hashomer, Helsinki committee approval number: SMC-7414-09

### Mice

The following mouse strains were used: C57BL/6J (wild-type), C57BL/6.SJL (PtprcaPep3b; Ly5.1), B6.129S7-Rag1tm1Mom/J (Rag1^-/-^), C57BL/6-Tg(TcraTcrb)1100Mjb/J (OT1), Slfn2-elektra, B6.Cg-Thy1a/Cy Tg(TcraTcrb)8Rest/J (Pmel), B6;129S-Gt(ROSA)26Sortm1(CAG-COX8A/Dendra2)Dcc/J (mito-Dendra), Mice were maintained under SPF conditions according to FELASA recommendations in the animal facilities of the The Hebrew University of Jerusalem and, the Biomedical Research Center, University of Marburg. All studies involving mice were done in accordance with institutional regulations governing animal care and use. NIH approval number: OPRR-A01-501. Mice were with C57BL/6 background and 8-12 weeks old mice used.

### Isolation of hyperpolarized and depolarized naïve CD8^+^ T cells

CD8^+^ T cells from spleens were negatively enriched using an EasySep™ Mouse CD8^+^ T Cell Isolation Kit (#19853) according to the manufacturer’s instructions (STEMCELL Technologies). Enriched CD8^+^ T cells (92% - 96% purity) were then stained with TMRM, a mix of anti-CD44, and anti-CD62L antibodies. Cells were then FACS-sorted for CD44-, CD62L^+^, TMRM high or TMRM low (99.5% purity) using a FACSAria (BD).

### Listeria infection

Ly5.1 mice were infected with 1×10^5^ CFU LmOVA via intraperitoneal route. dpT and hpT cells were sorted from OTI-Ly5.2 transgenic mice. 2×10^4^ or 8×10^4^ of the sorted cells were then intraperitoneally transferred into the sex- and age-matched Ly5.1 recipients that were infected with LmOVA. On day 8 post transfer, OT-I cell response in the spleen were analyzed by flow cytometry.

### Flow cytometry and ImageStream analyses

All antibodies were purchased from Biolegend, eBioscience or BD Bioscience. For TMRM staining, 10^6^ cells were suspended in 50nM TMRM in PBS (w/o Ca^2+^ and Mg^2+^) containing 1% FBS (w/o EDTA) and incubated for 30 minutes at 37°C following a washing step with PBS (w/o Ca^2+^ and Mg^2+)^ containing 1% FBS (w/o EDTA). The relevant antibody mixture diluted in PBS containing 1% FBS. Cells were then analyzed using a Gallios flow cytometer (Beckman Coulter) or Amnis® ImageStreamX Mark II imaging flow cytometer. Analysis was performed by FCS-express, FlowJo or the IDEAS software.

Antibodies for flow cytometry included anti-CD4-Pacific Blue (clone: GK1.5; BioLegend); anti–CD8α-PerCP/Cy5.5 & Violet 500 (clone: 53–6.7; BioLegend & BD Bioscience); anti-CD44-Alexa Fluor 700 & APC/Cy7 (clone: IM7; BioLegend), anti-CD45.1-APC (clone: A20; BioLegend); anti-CD45.2-Alexa Fluor 700 & BV510 (clone: 104; eBioscience & BioLegend); anti-CD62L-eVolve 605 & FITC (clone: MEL-14; eBioscience; anti-TCRβ-FITC (clone: H57-597; BD Bioscience); anti-CD69-APC (clone: H1.2F3; BioLegend); anti-IFNγ-PerCP/Cy5.5 (clone: XMG1.2; BioLegend); anti-TNF35-FITC (clone: MP6-XT22; eBioscience); anti-IL-2-PE (clone: JES6-5H4; eBioscience); anti-Eomes-FITC (clone: Dan11mag; eBioscience); anti-T-bet-PE/Cy7 (clone: 4B10; BioLegend).

### Adoptive transfer

Isolated hpT or dpT or total naïve CD8^+^ T cells were stained with CellTrace violet (1µM) and transferred intraperitoneally to recipient mice. At 7 - 10 days after adoptive transfer, blood cells or splenocytes were prepared, stained with TMRM and anti CD45.1, CD45.2 and CD8 antibodies. Then stained cells were analyzed by flow cytometry for CellTrace dilution. Donor cells were obtained from either B6;129S-Gt(ROSA)26Sortm1(CAG-COX8A/Dendra2)Dcc/J, C57BL/6.SJL (PtprcaPep3b; Ly5.1) or C57BL/6-Tg(TcraTcrb)1100Mjb/J mice. C57BL6/J (Ly5.2), (Ly5.2/ Ly5.1), Elektra or Rag1^-/-^ mice were used as recipients.

### *In vitro* T cell proliferation and activation

Sorted hpT, dpT, total naïve CD8^+^ or splenocytes were stained with CellTrace Violet (1μM), activated with SIINFEKL (63ng/ml - 500ng/ml) or plate-bound anti-CD3 (1μg ml, clone 145-2C11, Biolegend) and anti-CD28 (1μg/ml, clone 37.51, Biolegend) and analyzed by flow-cytometry every 24 hours for 3 or 4 days as indicated.

Sorted hpT, dpT and SP8 thymocytes (CD8^+^, TCRβ^+^, CD69^-^) from OT-I mice were stimulated with plate-bound anti-CD3 mAb (5µg/ml, clone 145-2C11, Biolegend), and soluble anti-CD28 (1µg/ml, clone 37.51, Biolegend) in the presence of rhIL-2 (50U/ml, Novartis) and anti-IFNγ (5µg/ml, clone XMG1.2, Biolegend). For intracellular cytokine staining, murine cells were restimulated after 72h of culture with 10nM SIINFEKL_257-264_ in the presence of Brefeldin A (5µg/ml; Biolegend) for 4h. After fixation with 2% para-formaldehyde, intracellular staining in Saponin Buffer (0.1% Saponin, 1% BSA in PBS) was done using anti-IFNγ (clone XMG1.2, Biolegend), anti-TNF (clone: MP6-XT22; eBioscience); anti-IL-2 (clone: JES6-5H4; eBioscience) for 45min at 4°C. Staining of transcription factors was performed after 36 h without restimulation using the FOXP3/Fixation-Kit (eBioscience, 00-5521-00). For intracellular staining, anti-Eomes (clone: Dan11mag; eBioscience); anti-T-bet (clone: 4B10; BioLegend); antibodies were diluted in 0.1% Saponin in PBS. The cells were acquired on the Attune NxT Cytometer (Thermofisher Scientific).

For the experiments with Sinefungin the cells were pre-incubated with 30µM Sinefungin for 3h and then seeded into plates coated with anti-CD3 as described above with addition of soluble anti-CD28, rhIL-2, anti-IFNγ and rmIL-12 (2ng/ml, Peprotech). Intracellular levels of T-bet, IFNγ and TNF were analyzed after 36 h of culture.

### *In vitro* T cell S35-Met Cytotoxicity assay

The ability of CD8^+^ T cells to kill target cells was directly measured using the standard S35-Met release assays. Sorted hpT and dpT cells were activated with anti-CD3 (0.05 μg/well) and anti-CD28 (0.05 μg/well). The target cells (P815) were labeled with [S35]-Methionine 12 hours prior to the assay. Labeled targets (5000 cells/well) were added with different amounts of activated CD8^+^ T cells (various E:T ratios) and then diluted with RPMI medium in 96-U shaped plates at 37ºC for 5 h. Following incubation, plates were centrifuged (1600rpm, 5min, 4ºC) and supernatants (50µl) were collected and transferred to opaque Opti-plates (Packard). Wells were added with 150µl scintillation liquid (Packard) and analyzed by a β-counter (Packard). The total amount of labeled cells was determined by adding 100μl of 0.1M NaOH to an equal amount of targets (5000/well) that were not added with effectors. Spontaneous release was determined by measuring radioactive readings (as above) in wells that were given identical treatment as the experimental wells, but were not added with CD8^+^ T cells. Final specific lysis was calculated as follows: ((radioactive reading – spontaneous release) / (total labeling – spontaneous release))*100 = specific lysis.

### T cells depletion assay

T cells were depleted by injecting i.p. to 8-week-old mice, with anti-Thy1.2 antibody. Eight days post antibody injection, the polarization levels of the newly emerging CD8^+^ T cells in the spleen of these mice were measured by TMRM staining using flowcytometry analysis.

### Metabolic assays

OCR was measured using a 24-well XF extracellular flux analyzer (EFA) (Seahorse Bioscience). Isolated hpT and dpT cells (1×10^6^ cells per well) were seeded in Seahorse XF24 designated plates using Cell-Tak (Corning) adherent and assayed according to manufacturer instructions. ATP was measured using ATP-lite assay kit (Perkin Elmer) using (5×10^5^ cells/well and 1×10^6^ cells/well). Lactate quantification was performed using Picoprobe Lactate assay kit (abcam); hpT and dpT cells were isolated and incubated (1×10^6^ cells/well) for 3 h in PBS with 1% BSA with or without 25 mM Glucose, and assayed according to manufacturer instructions.

### Targeted metabolic analysis

Isolated hpT and dpT cells were cultured in 96 well plate (1× 10^6^ cells/well), suspended in RPMI supplemented with 10% dialyzed Fetal Bovine Serum and 100μM Alanine with or without 30µM Sinefungin. Following 5 or 24 h, were then extracted for metabolomics LC-MS analysis.

Medium extracts: Twenty microliters of culture medium was added to 980 μl of a cold extraction solution (−20°C) composed of methanol, acetonitrile, and water (5:3:2). Cell extracts: Cells were rapidly washed 3 times with ice-cold PBS, after which intracellular metabolites were extracted with 100μl of ice-cold extraction solution for 5 min at 4°C. Medium and cell extracts were centrifuged (10 min at 16,000g) to remove insoluble material, and the supernatant was collected for LC-MS analysis. Metabolomics data was normalized to protein concentrations using a modified Lowry protein assay.

LC-MS metabolomics analysis was performed as described previously(MacKay et al., 2015). Briefly, Thermo Ultimate 3000 high-performance liquid chromatography (HPLC) system coupled to Q-Exactive Orbitrap Mass Spectrometer (Thermo Fisher Scientific) was used with a resolution of 35,000 at 200 mass/charge ratio (m/z), electrospray ionization, and polarity switching mode to enable both positive and negative ions across a mass range of 67 to 1000 m/z. HPLC setup consisted ZIC-pHILIC column (SeQuant; 150 mm x 2.1 mm, 5 μm; Merck), with a ZIC-pHILIC guard column (SeQuant; 20 mm x 2.1 mm). 5 μl of Biological extracts were injected and the compounds were separated with mobile phase gradient of 15 min, starting at 20% aqueous (20 mM ammonium carbonate adjusted to pH.2 with 0.1% of 25% ammonium hydroxide) and 80% organic (acetonitrile) and terminated with 20% acetonitrile. Flow rate and column temperature were maintained at 0.2 ml/min and 45°C, respectively, for a total run time of 27 min. All metabolites were detected using mass accuracy below 5 ppm. Thermo Xcalibur was used for data acquisition. TraceFinder 4.1 was used for analysis. Peak areas of metabolites were determined by using the exact mass of the singly charged ions. The retention time of metabolites was predetermined on the pHILIC column by analyzing an in-house mass spectrometry metabolite library that was built by running commercially available standards.

### ATAC-seq sample preparation and sequencing

ATAC-seq. samples were prepared as previously described(Buenrostro et al., 2013) with minor modifications. In brief, 5×10^4^ dpT and hpT cells were sorted from spleen and peripheral draining lymph nodes. Cells were lysed by 50µl ATAC-lysis buffer (10mM Tris-HCl pH7.4, 10mM NaCl, 3mM MgCl2, 0.1% NP40, 0.1% Tween-20 and 0.01% digitonin) for 3 min on ice. Resuspend cells in 1ml cold ATAC resuspension buffer (10mM Tris-HCl pH7.4, 10mM NaCl, 3mM MgCl2) supplied with 0.1% Tween 20 and centrifuge for 10min at 4°C, 500xg. Cell pellet was resuspended in 50µl transposase reaction mix (25µl 2xTD buffer, 2.5µl Transposase, 16.5µl 1x PBS, 5µl nuclease-free water, 0.5µl 1% digitonin, 0.5µl 10% Tween 20) and incubated on a shaker at 1000rpm for 30min at 37°C. DNA was purified with Zymo clean and concentrator-5 kit. 1 µl of the eluted DNA was used in a quantitative 10 µL PCR reaction (1.25 µM forward and reverse custom Nextera primers, 1x SYBR green final concentration) to estimate the optimum number of amplification cycles with the following program 72°C 5 min; 98°C 30 s; 25 cycles: 98°C 10 s, 63°C 30 s, 72°C 1 min; the final amplification of the library was carried out using the same PCR program and the number of cycles according to the Cq value of the qPCR. Library amplification using custom Nextera primers was followed by SPRI size selection with AmpureXP beads to exclude fragments larger than 1500 bp and smaller than 150 bp. DNA concentration was measured by Qubit fluorometer (Life Technologies). The libraries were sequenced using the Illumina NextSeq 550 system.

### Processing of ATAC-seq data

ATAC-seq samples were aligned to the GRCm38 reference genome with Bowtie2 (with the setting “--very-sensitive”). Quality control for the samples was performed with the ATACseqQC (v1.12.3) (Ou et al., 2018) and ChIPQC (v1.24.1) packages. Peaks on the individual samples were called with MACS2(Zhang et al., 2008), with a FDR q-value of 0.1 and parameters “--nomodel --shift -100 --extsize 200”. A consensus peak list was created by combining all peaks in all samples, excluding the ENCODE blacklisted regions to avoid artifacts. The numbers of reads in each consensus region across all samples were computed with Rsubread’s routine featureCounts (Liao et al., 2014). Annotation of the consensus regions to the respective genes (based on genomic proximity) was carried out with ChIPseeker (Yu et al., 2015). Exploratory data analysis (including Principal Component Analysis) was performed with pcaExplorer [Marini & Binder, v2.15.2](Marini and Binder, 2019). Regions with differential ATAC-seq signal were identified using DESeq2 (v1.29.6)(Love et al., 2014) for normalization, variance estimation, and testing, using a FDR cutoff of 0.1. Functional enrichment analysis was performed providing the lists of genes that the differentially occupied regions were assigned to, with functions from the clusterProfiler package (G. Yu et al., 2012). Normalized bigwig tracks were generated with deepTools (Ramírez et al., 2014). Heatmaps were generated with the variance stabilizing transformed values for the normalized counts of the selected regions, standardized by row.

### Statistical analysis

The statistical significance of differences was determined by ANOVA. Differences with a P value of less than 0.05 were considered statistically significant. Graph prism and Perseus programs were use. MS data was normalized by ranking, when applicable, non-values were plugged with replicates mean to prevent zeros bias.

### Protein mass spectrometry

Sample preparation - samples, frozen at -20°C were subjected to tryptic digestion, performed in the presence of 0.05% ProteaseMAX Surfactant (from Promega Corp., Madison, WI, USA). The peptides were then desalted on C18 Stage tips (Rappsilber et al., 2007). A total of 0.5 µg of peptides were injected into the mass spectrometer.

LC MS/MS analysis - MS analysis was performed using a Q Exactive Plus mass spectrometer (Thermo Fisher Scientific) coupled on-line to a nanoflow UHPLC instrument (Ultimate 3000 Dionex, Thermo Fisher Scientific). Eluted peptides were separated over a 60-min gradient run at a flow rate of 0.3 μl/min on a reverse phase 25-cm-long C18 column (75um ID, 2µm, 100Å, Thermo PepMap®RSLC). The survey scans (380–2,000 m/z, target value 3E6 charges, maximum ion injection times 50 ms) were acquired and followed by higher energy collisional dissociation (HCD) based fragmentation (normalized collision energy 285). A resolution of 70,000 was used for survey scans and up to 15 dynamically chosen most abundant precursor ions were fragmented (isolation window 1.6 m/z). The MS/MS scans were acquired at a resolution of 17,500 (target value 5E4 charges, maximum ion injection times 57 ms). Dynamic exclusion was 60 sec.

MS data analysis - Mass spectra data were processed using the MaxQuant computational platform, version 1.5.3.12. Peak lists were searched against the Homo sapiens Uniprot FASTA sequence database containing a total of 26,199 reviewed entries or a custom FATSA file containing mitocarta genes. Peptides with minimum of seven amino-acid length were considered and the required FDR was set to 1% at the peptide and protein level. Protein identification required at least 3 unique or razor peptides per protein group. The dependent-peptide and match-between-runs options were used.

## Supplemental information

Supplemental information includes 5 figures, 4 Tables

## Author contributions

A.S., M.B. and MH designed and performed research, analyzed data and wrote the manuscript; X.Z., I.A. L.C., L.Z., H.R., I.O., A.S., M.K., A.D., F.M., O.T., R.S., C.S., R.C.H., E.G., performed research.

## Acknowledgments

The authors thank Dr. Ofer Mandelboim (The Hebrew University), Dr. David Chan (Caltech) and Dr. Atan Gross (Weizmann Institute) for critical reviews, Dr. Tsvee Lapidot (Weizmann Institute) for mito-Dendra2 mice, Dr. Michal Baniyash (The Hebrew University) for PMEL17 mice, Dr. Avihai Hovav (The Hebrew University) for SIINFEKL peptide, and Dr. Eli Pikarsky (The Hebrew University) for the anti-Thy1 antibody.

This work was supported by grants from the Israel Science Foundation grant No. 1596/17, German-Israeli Foundation grant No. I-224-414.11-2017, Israel Cancer Research Foundation (ICRF), and by the Memorial Sloan Kettering Cancer Center Core Grant (P30 CA008748).

## Supplementary Figures

**Figure S1.**
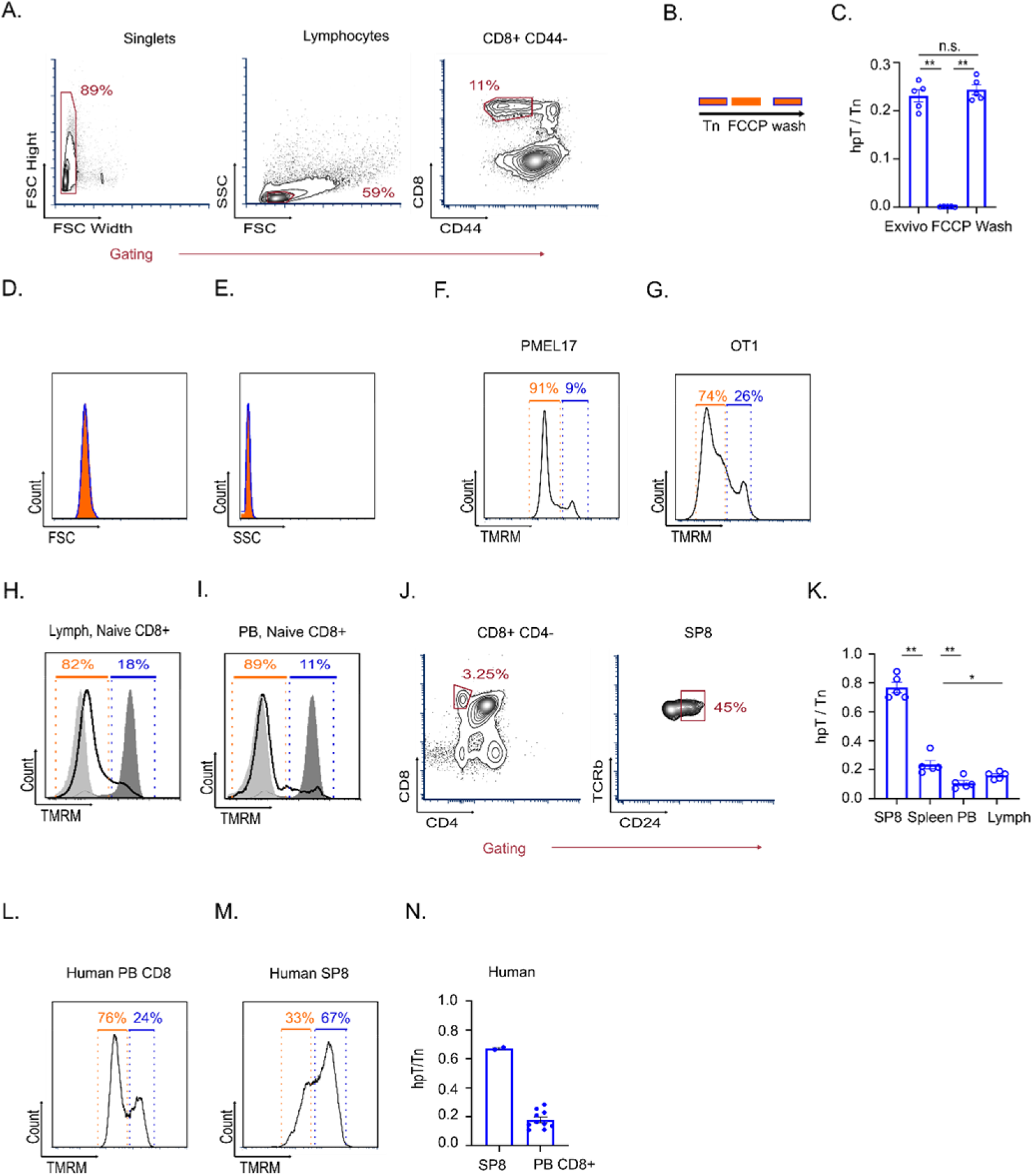
Mitochondrial polarization staining distinguishes between two developmental stages of naïve CD8^+^ T cell: **A**. Gating strategy for identification of Tn from mouse spleen **B**. Schematics of experiment presented in C. **C**. Mouse splenocytes were treated with FCCP for 30 minutes. Then cells were divided into two different groups; 1) continued to grow in the same FCCP-containing medium, or 2) medium containing FCCP was washed, and cells were incubated with medium without FCCP. Thirty minutes after incubation cells stained with TMRM and anti-CD8 antibody and subjected to flow cytometry analysis. Splenocytes were incubated for 1 hour in a normal medium as a control. Bar graph, quantification of relative abundance of hpT with respect to total Tn in the control (*ex vivo*), FCCP and washed cells. **D**. Representative overlay histogram showing the FSC signal gated on dpT (orange) and hpT (blue). **E**. Representative overlay histogram showing the SSC signal gated on dpT (orange) and hpT (blue). **F**. Representative histogram showing TMRM intensity in PMEL17 mouse splenocytes gated on CD8^+^ CD44^low^ cells. dpT (orange region) and hpT (blue region). **G**. Same as in F., in OT-1 mouse. **H**. Overlay histogram showing TMRM intensity, gated on CD8^+^ CD44^low^ T cells from mouse lymph nodes and treated with either FCCP (minimal TMRM staining-light gray), oligomycin (maximal TMRM staining-dark gray) or left untreated (black line). dpT (orange region) hpT (blue region) **I**. Same as in H, from peripheral blood (PB). **J**. Gating strategy for identification of SP8 from mouse thymus. **K**. Bar graph, summary of hpT relative abundance in samples from different tissues (n=5 mice). **L**. Representative histogram showing TMRM intensity in human peripheral blood gated on CD8^+^ CD45RA^+^ CD62L^+^ cells. dpT (orange region) and hpT (blue region). **M**. Representative histogram showing TMRM intensity in human Thymus gated on CD8^+^ CD4^-^ cells. dpT (orange region) and hpT (blue region). **N**. summary of experiments presented in L and M (PB n=10, Thymus n=2). **Statistics** | Mann-Whitney test - P-values – C. **=0.0079, K. *=0.0159 **=0.0072 | error bars represent s.e.m

**Figure S2.**
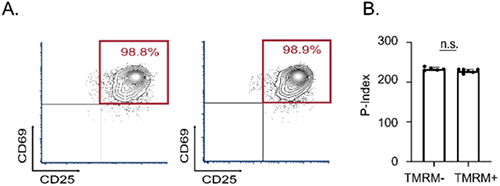
Immature-hpT are hyporesponsive to diverse stimuli in respect to mature-dpT both *in vitro* and *in vivo*: (A-B) Cell Trace labeled mouse splenocytes were activated *in vitro* in the presence or absence of 1 µM TMRM. **A**. Representative scatter plot showing CD25 and CD69 intensities gated on CD8^+^ T cells 24 hours following activation. **B**. Bar graph showing the proliferation index, assessed by dilution of Cell Trace intensity, of CD8^+^ T cells 72 hours post stimuli (n=6 biological replicates). **Statistics** | Mann-Whitney test - P-values – B. n.s. error bars represent s.e.m

**Figure S3.**
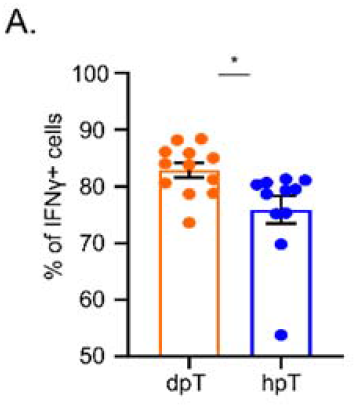
Immature-hpT maintain distinct metabolic profile in respect to mature-dpT: hpT or dpT cells isolated from syngeneic OT-1 Ly5.2 mice were adoptively transferred at 1*10^5 cells/mouse into LmOVA infected Ly5.1 recipients. Eight days post transfer activated CD8^+^ CD45.2^+^ cells were permeabilized and analyzed by flow cytometry for expression levels of IFNγ. **A**. Bar graph, summary of relative abundance of IFNγ^+^ cells dpT (orange) hpT (blue), (n=12 mice). **Statistics** | Mann-Whitney test - P-values – A. *=0.0179 | error bars represent s.e.m

**Figure S4.**
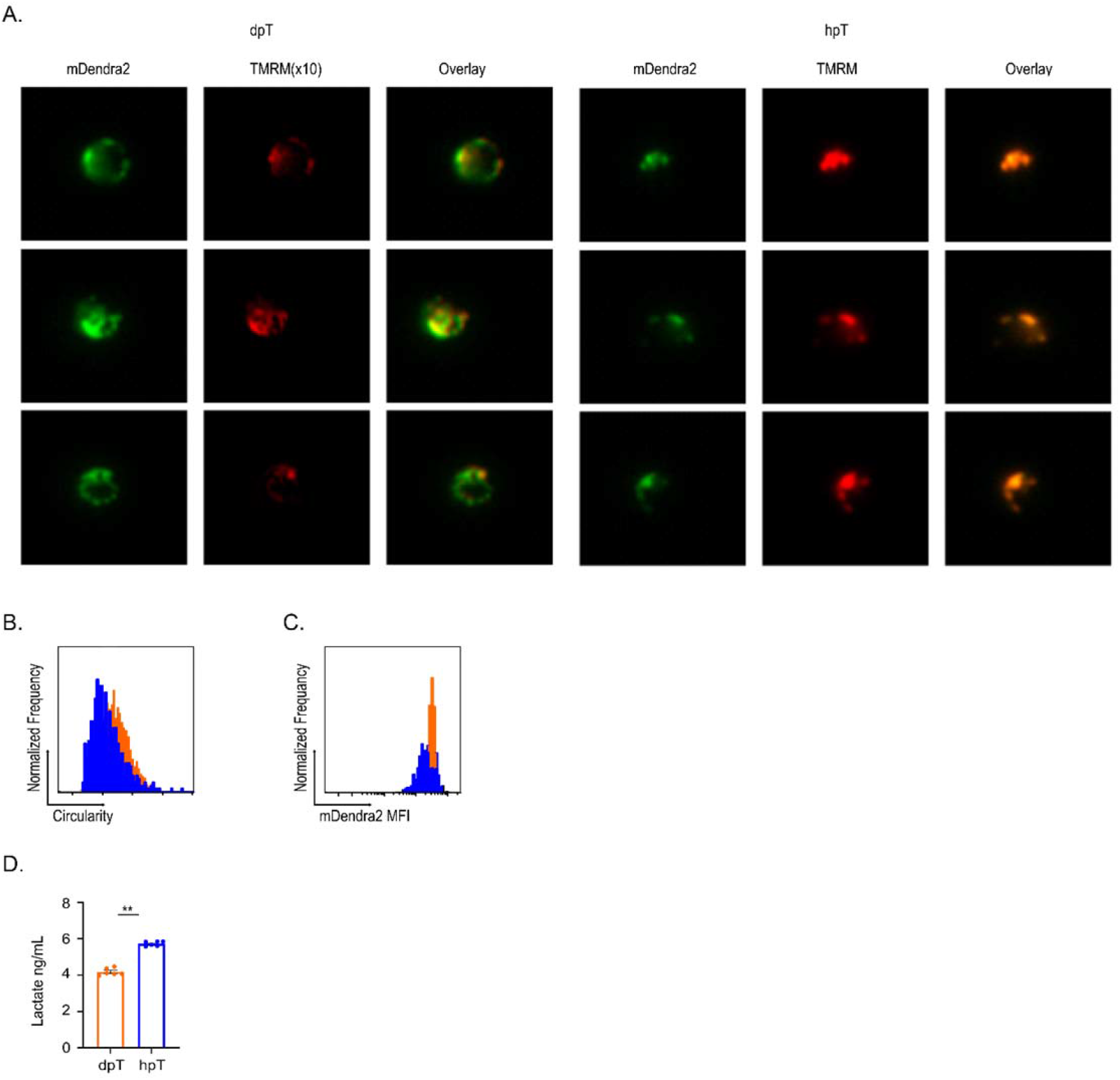
hpT maintain distinct metabolic profile in respect to dpT: **A**. Representative ImageStream images of mDendra2 (green) as a marker of total mitochondrial mass and TMRM staining (red) of hpT (right three panels) and dpT (left three panels) cells from spleens of mtDenedra2 mice (n=3 biological replicates). **B**. ImageStream overlay histogram of mtDendra2 circularity, representing the cellular distribution of mtDendra2 in hpT (bllue) and dpT (orange) cells **C**. ImageStream overlay histogram, Log mean intensity of mtDendra2 in hpT (blue) and dpT (orange) cells. **D**. Isolated dpT and hpT were incubated for 2 hours in PBS containing 25mM glucose. Lactate concentration was then quantified by colorimetric assay. dpT (orange), hpT (blue). (n=6 biological replicates) **Statistics** | Mann-Whitney test - P-values – D. **=0.0022 | error bars represent s.e.m

**Figure S5.**
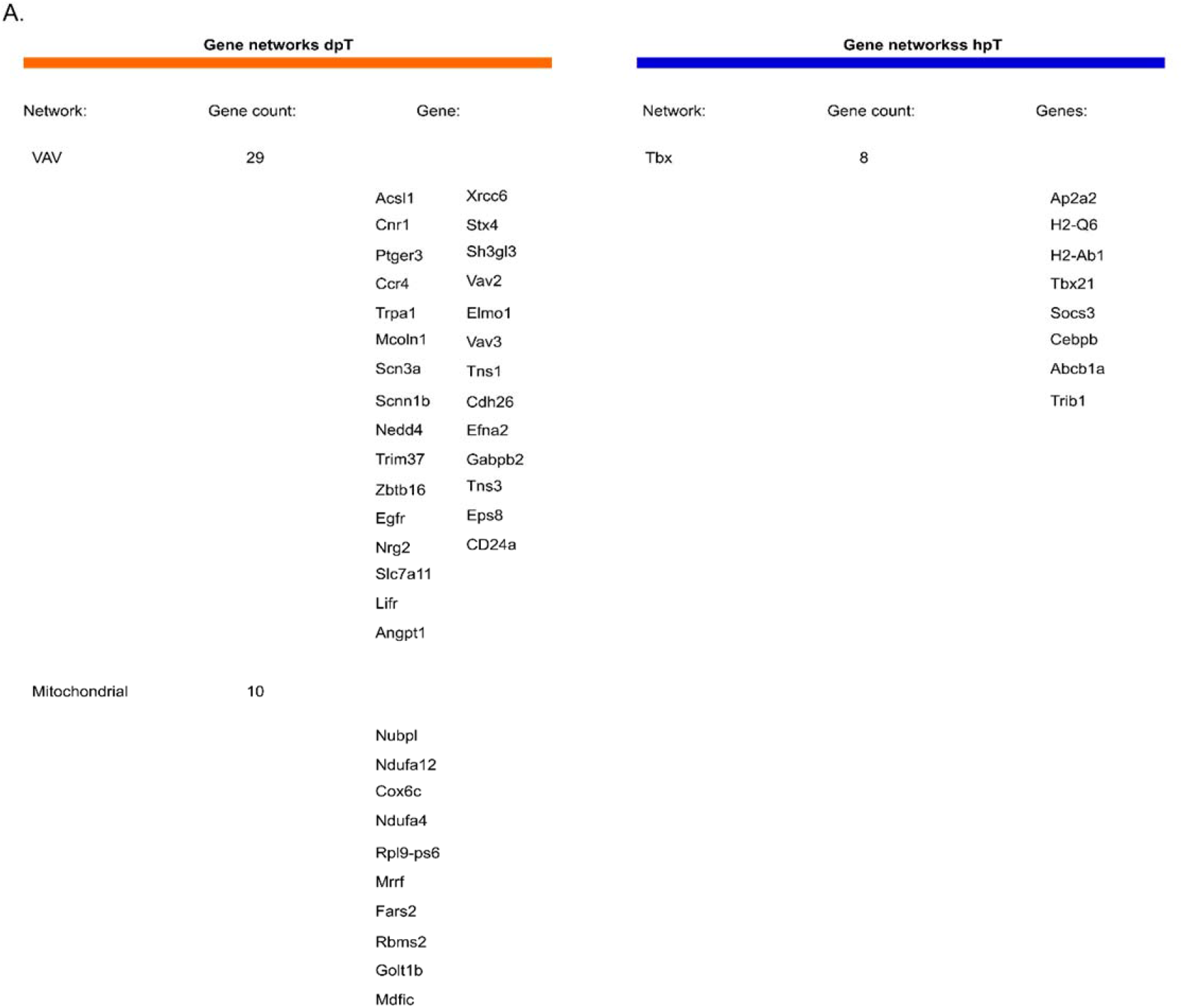
Reduced chromatin accessibility at Tbx21 locus correlates with decreased T-BET expression in immature-hpT cells: **A**. Detailed table listing individual genes in each string network presented in Figure 5.

## References

Abdelsamed HA, Zebley CC, Nguyen H, Rutishauser RL, Fan Y, Ghoneim HE, Crawford JC, Alfei F, Alli S, Ribeiro SP, Castellaw AH, McGargill MA, Jin H, Boi SK, Speake C, Serti E, Turka LA, Busch ME, Stone M, Deeks SG, Sekaly RP, Zehn D, James EA, Nepom GT, Youngblood B. 2020. Beta cell-specific CD8+ T cells maintain stem cell memory-associated epigenetic programs during type 1 diabetes. Nature Immunology 21:578–587. doi:10.1038/s41590-020-0633-5

Bailis W, Shyer JA, Zhao J, Carlos J, Canaveras G, Khazal J Al, Qu R, Steach HR, Bielecki P, Khan O, Kluger Y, Maher LJ, Rabinowitz J, Craft J. 2020. differentiation and function. Nature 571:403–407. doi:10.1038/s41586-019-1311-3.Distinct

Bantug GR, Galluzzi L, Kroemer G, Hess C. 2017. The spectrum of T cell metabolism in health and disease. Nature Reviews Immunology. doi:10.1038/nri.2017.99

Berberich-Siebelt F, Klein-Hessling S, Hepping N, Santner-Nanan B, Lindemann D, Schimpl A, Berberich I, Serfling E. 2000. C/EBPβ enhances IL-4 but impairs IL-2 IFN-γ induction in T cells. European Journal of Immunology 30:2576–2585. doi:10.1002/1521-4141(200009)30:9<2576::AID-IMMU2576>3.0.CO;2-N

Berger M, Krebs P, Crozat K, Li X, Croker BA, Siggs OM, Popkin D, Du X, Lawson BR, Theofilopoulos AN, Xia Y, Khovananth K, Moresco EMY, Satoh T, Takeuchi O, Akira S, Beutler B. 2010. An Slfn2 mutation causes lymphoid and myeloid immunodeficiency due to loss of immune cell quiescence. Nature Immunology 11:335–343. doi:10.1038/ni.1847

Berkley AM, Fink PJ. 2014a. Cutting Edge: CD8 + Recent Thymic Emigrants Exhibit Increased Responses to Low-Affinity Ligands and Improved Access to Peripheral Sites of Inflammation. The Journal of Immunology. doi:10.4049/jimmunol.1401870

Berkley AM, Fink PJ. 2014b. Cutting Edge: CD8 + Recent Thymic Emigrants Exhibit Increased Responses to Low-Affinity Ligands and Improved Access to Peripheral Sites of Inflammation. The Journal of Immunology 193:3262–3266. doi:10.4049/jimmunol.1401870

Boursalian TE, Golob J, Soper DM, Cooper CJ, Fink PJ. 2004. Continued maturation of thymic emigrants in the periphery. Nature Immunology. doi:10.1038/ni1049

Buck MDD, O’Sullivan D, Klein Geltink RII, Curtis JDD, Chang CH, Sanin DEE, Qiu J, Kretz O, Braas D, van der Windt GJJW, Chen Q, Huang SCC, O’Neill CMM, Edelson BTT, Pearce EJJ, Sesaki H, Huber TBB, Rambold ASS, Pearce ELL. 2016. Mitochondrial Dynamics Controls T Cell Fate through Metabolic Programming. Cell 166:63–76. doi:10.1016/j.cell.2016.05.035

Buenrostro JD, Giresi PG, Zaba LC, Chang HY, Greenleaf WJ. 2013. Transposition of native chromatin for fast and sensitive epigenomic profiling of open chromatin, DNA-binding proteins and nucleosome position. Nature Methods 10:1213–1218. doi:10.1038/nmeth.2688

Carr EL, Kelman A, Wu GS, Gopaul R, Senkevitch E, Aghvanyan A, Turay AM, Frauwirth KA. 2010. Glutamine Uptake and Metabolism Are Coordinately Regulated by ERK/MAPK during T Lymphocyte Activation. The Journal of Immunology 185:1037–1044. doi:10.4049/jimmunol.0903586

Chapman NM, Boothby MR, Chi H. 2020. Metabolic coordination of T cell quiescence and activation. Nature Reviews Immunology 20:55–70. doi:10.1038/s41577-019-0203-y

Cunningham CA, Bergsbaken T, Fink PJ. 2017. Cutting Edge: Defective Aerobic Glycolysis Defines the Distinct Effector Function in Antigen-Activated CD8 + Recent Thymic Emigrants. The Journal of Immunology 198:4575–4580. doi:10.4049/jimmunol.1700465

Cunningham CA, Helm EY, Fink PJ. 2018a. Reinterpreting recent thymic emigrant function: defective or adaptive? Current Opinion in Immunology 51:1–6. doi:10.1016/j.coi.2017.12.006

Cunningham CA, Helm EY, Fink PJ. 2018b. Reinterpreting recent thymic emigrant function: defective or adaptive? Current Opinion in Immunology. doi:10.1016/j.coi.2017.12.006

Cunningham CA, Hoppins S, Fink PJ. 2018c. Cutting Edge: Glycolytic Metabolism and Mitochondrial Metabolism Are Uncoupled in Antigen-Activated CD8 + Recent Thymic Emigrants. The Journal of Immunology 201:1627–1632. doi:10.4049/jimmunol.1800705

Duckworth BC, Lafouresse F, Wimmer VC, Broomfield BJ, Dalit L, Alexandre YO, Sheikh AA, Qin RZ, Alvarado C, Mielke LA, Pellegrini M, Mueller SN, Boudier T, Rogers KL, Groom JR. 2021. Effector and stem-like memory cell fates are imprinted in distinct lymph node niches directed by CXCR3 ligands. Nature Immunology 22:434–448. doi:10.1038/s41590-021-00878-5

Fink PJ. 2013. The biology of recent thymic emigrants. Annual Review of Immunology 31:31–50. doi:10.1146/annurev-immunol-032712-100010

Fink PJ, Hendricks DW. 2011. Post-thymic maturation: Young T cells assert their individuality. Nature Reviews Immunology. doi:10.1038/nri3028

Fox CJ, Hammerman PS, Thompson CB. 2005. Fuel feeds function: Energy metabolism and the T-cell response. Nature Reviews Immunology 5:844–852. doi:10.1038/nri1710

Frauwirth KA, Riley JL, Harris MH, Parry R V, Rathmell JC, Plas DR, Elstrom RL, June CH, Thompson CB. 2002. The CD28 Signaling Pathway Regulates Glucose Metabolism of metabolism in response to changes in cellular conditions. However, it has recently been shown that signals from cell surface receptors are required to control the ability of resting cells to tak. Immunity 16:769–777.

Friesen TJ, Ji Q, Fink PJ. 2016a. Recent thymic emigrants are tolerized in the absence of inflammation. The Journal of Experimental Medicine. doi:10.1084/jem.20151990

Friesen TJ, Ji Q, Fink PJ. 2016b. Recent thymic emigrants are tolerized in the absence of inflammation. Journal of Experimental Medicine 213:913–920. doi:10.1084/jem.20151990

George AJT, Ritter MA. 1996. Thymic involution with ageing: Obsolescence or good housekeeping? Immunology Today 17:267–272. doi:10.1016/0167-5699(96)80543-3

Gerriets VA, Rathmell JC. 2012. Metabolic pathways in T cell fate and function. Trends in Immunology 33:168–173. doi:10.1016/j.it.2012.01.010

Goll MG, Bestor TH. 2005. Eukaryotic cytosine methyltransferases. Annual Review of Biochemistry 74:481–514. doi:10.1146/annurev.biochem.74.010904.153721

Gros C, Chauvigné L, Poulet A, Menon Y, Ausseil F, Dufau I, Arimondo PB. 2013. Development of a universal radioactive DNA methyltransferase inhibition test for highthroughput screening and mechanistic studies. Nucleic Acids Research 41. doi:10.1093/nar/gkt753

Han BY, Wu S, Foo CS, Horton RM, Jenne CN, Watson SR, Whittle B, Goodnow CC, Cyster JG. 2014. Zinc finger protein Zfp335 is required for the formation of the naïve T cell compartment. eLife 3:1–28. doi:10.7554/eLife.03549

Herek TA, Bouska A, Lone WG, Heavican TB, Amador C, Sharma S, Francesco d’Amore, Chan WC, Iqbal J. 2020. DNMT3A Mutations Identify a Prognostic Subgroup in Peripheral T-Cell Lymphoma. Blood 136:38–39. doi:10.1182/blood-2020-143335

Ho PC, Bihuniak JD, MacIntyre AN, Staron M, Liu X, Amezquita R, Tsui YC, Cui G, Micevic G, Perales JC, Kleinstein SH, Abel ED, Insogna KL, Feske S, Locasale JW, Bosenberg MW, Rathmell JC, Kaech SM. 2015. Phosphoenolpyruvate Is a Metabolic Checkpoint of Anti-tumor T Cell Responses. Cell 162:1217–1228. doi:10.1016/j.cell.2015.08.012

Hogquist KA, Xing Y, Hsu F-C, Shapiro VS. 2015. T Cell Adolescence: Maturation Events Beyond Positive Selection. The Journal of Immunology 195:1351–1357. doi:10.4049/jimmunol.1501050

Hooftman A, Angiari S, Hester S, Corcoran SE, Runtsch MC, Ling C, Ruzek MC, Slivka PF, McGettrick AF, Banahan K, Hughes MM, Irvine AD, Fischer R, O’Neill LAJ. 2020. The Immunomodulatory Metabolite Itaconate Modifies NLRP3 and Inhibits Inflammasome Activation. Cell Metabolism 32:468-478.e7. doi:10.1016/j.cmet.2020.07.016

Hsu F-C, Belmonte PJ, Constans MM, Chen MW, McWilliams DC, Hiebert SW, Shapiro VS. 2015. Histone Deacetylase 3 Is Required for T Cell Maturation. The Journal of Immunology 195:1578–1590. doi:10.4049/jimmunol.1500435

Hsu F-C, Pajerowski AG, Nelson-Holte M, Sundsbak R, Shapiro VS. 2011. NKAP is required for T cell maturation and acquisition of functional competency. The Journal of Experimental Medicine. doi:10.1084/jem.20101874

Jacobs SR, Herman CE, MacIver NJ, Wofford JA, Wieman HL, Hammen JJ, Rathmell JC. 2008. Glucose Uptake Is Limiting in T Cell Activation and Requires CD28-Mediated Akt-Dependent and Independent Pathways. The Journal of Immunology 180:4476–4486. doi:10.4049/jimmunol.180.7.4476

Joshi NS, Cui W, Chandele A, Lee HK, Urso DR, Hagman J, Gapin L, Kaech SM. 2007. Inflammation Directs Memory Precursor and Short-Lived Effector CD8+ T Cell Fates via the Graded Expression of T-bet Transcription Factor. Immunity 27:281–295. doi:10.1016/j.immuni.2007.07.010

Kaech SM, Cui W. 2012. Transcriptional control of effector and memory CD8+ T cell differentiation. Nature Reviews Immunology 12:749–761. doi:10.1038/nri3307

Katajisto P, Döhla J, Chaffer CL, Pentinmikko N, Marjanovic N, Iqbal S, Zoncu R, Chen W, Weinberg RA, Sabatini DM. 2015. Asymmetric apportioning of aged mitochondria between daughter cells is required for stemness. Science. doi:10.1126/science.1260384

Kim HK, Waickman AT, Castro E, Flomerfelt FA, Hawk N V., Kapoor V, Telford WG, Gress RE. 2016. Distinct IL-7 signaling in recent thymic emigrants versus mature naïve T cells controls T-cell homeostasis. European Journal of Immunology 46:1669–1680. doi:10.1002/eji.201546214

Ladle BH, Li KP, Phillips MJ, Pucsek AB, Haile A, Powell JD, Jaffee EM, Hildeman DA, Gamper CJ. 2016. De novo DNA methylation by DNA methyltransferase 3a controls early effector CD8+ T-cell fate decisions following activation. Proceedings of the National Academy of Sciences of the United States of America 113:10631–10636. doi:10.1073/pnas.1524490113

Liao Y, Smyth GK, Shi W. 2014. FeatureCounts: An efficient general purpose program for assigning sequence reads to genomic features. Bioinformatics 30:923–930. doi:10.1093/bioinformatics/btt656

Love MI, Huber W, Anders S. 2014. Moderated estimation of fold change and dispersion for RNA-seq data with DESeq2. Genome Biology 15:1–21. doi:10.1186/s13059-014-0550-8

Lynch HE, Goldberg GL, Chidgey A, Van den Brink MRM, Boyd R, Sempowski GD. 2009. Thymic involution and immune reconstitution. Trends in Immunology 30:366–373. doi:10.1016/j.it.2009.04.003

Ma D, Wei Y, Liu F. 2013. Regulatory mechanisms of thymus and T cell development. Developmental and Comparative Immunology 39:91–102. doi:10.1016/j.dci.2011.12.013

MacIver NJ, Michalek RD, Rathmell JC. 2013. Metabolic regulation of T lymphocytes. Annu Rev Immunol. doi:10.1146/annurev-immunol-032712-095956

MacKay GM, Zheng L, Van Den Broek NJF, Gottlieb E. 2015. Analysis of Cell Metabolism Using LC-MS and Isotope TracersMethods in Enzymology. doi:10.1016/bs.mie.2015.05.016

Makaroff L. E., Hendricks DW, Niec RE, Fink PJ. 2009. Postthymic maturation influences the CD8 T cell response to antigen. Proceedings of the National Academy of Sciences. doi:10.1073/pnas.0812354106

Makaroff Lydia E., Hendricks DW, Niec RE, Fink PJ. 2009. Postthymic maturation influences the CD8 T cell response to antigen. Proceedings of the National Academy of Sciences of the United States of America 106:4799–4804. doi:10.1073/pnas.0812354106

Marini F, Binder H. 2019. PcaExplorer: An R/Bioconductor package for interacting with RNA-seq principal components. BMC Bioinformatics 20:1–8. doi:10.1186/s12859-019-2879-1

Miyajima M. 2020. Amino acids: Key sources for immunometabolites and immunotransmitters. International Immunology 32:435–446. doi:10.1093/intimm/dxaa019

Nakaya M, Xiao Y, Zhou X, Chang JH, Chang M, Cheng X, Blonska M, Lin X, Sun SC. 2014. Inflammatory T cell responses rely on amino acid transporter ASCT2 facilitation of glutamine uptake and mTORC1 kinase activation. Immunity 40:692–705. doi:10.1016/j.immuni.2014.04.007

Nastasi C, Willerlev-Olsen A, Dalhoff K, Ford SL, Gadsbøll ASØ, Buus TB, Gluud M, Danielsen M, Litman T, Bonefeld CM, Geisler C, Ødum N, Woetmann A. 2021. Inhibition of succinate dehydrogenase activity impairs human T cell activation and function. Scientific Reports 11:1–13. doi:10.1038/s41598-020-80933-7

Ou J, Liu H, Yu J, Kelliher MA, Castilla LH, Lawson ND, Zhu LJ. 2018. ATACseqQC: A Bioconductor package for post-alignment quality assessment of ATAC-seq data. BMC Genomics 19:1–13. doi:10.1186/s12864-018-4559-3

Pearce EL, Poffenberger MC, Chang CH, Jones RG. 2013. Fueling immunity: Insights into metabolism and lymphocyte function. Science 342. doi:10.1126/science.1242454

Pham AH, Mccaffery JM, Chan DC. 2012. Mouse lines with photo-activatable mitochondria to study mitochondrial dynamics. Genesis. doi:10.1002/dvg.22050

Pham D, Yu Q, Walline CC, Muthukrishnan R, Blum JS, Kaplan MH. 2013. Opposing Roles of STAT4 and Dnmt3a in Th1 Gene Regulation. The Journal of Immunology 191:902–911. doi:10.4049/jimmunol.1203229

Pietrocola F, Galluzzi L, Bravo-San Pedro JM, Madeo F, Kroemer G. 2015. Acetyl coenzyme A: A central metabolite and second messenger. Cell Metabolism 21:805–821. doi:10.1016/j.cmet.2015.05.014

Priyadharshini B, Welsh RM, Greiner DL, Gerstein RM, Brehm MA. 2010. Maturation-Dependent Licensing of Naive T Cells for Rapid TNF Production. PLoS ONE. doi:10.1371/journal.pone.0015038

Ramírez F, Dündar F, Diehl S, Grüning BA, Manke T. 2014. DeepTools: A flexible platform for exploring deep-sequencing data. Nucleic Acids Research 42:187–191. doi:10.1093/nar/gku365

Rappsilber J, Mann M, Ishihama Y. 2007. Protocol for micro-purification, enrichment, prefractionation and storage of peptides for proteomics using StageTips. Nature Protocols. doi:10.1038/nprot.2007.261

Rome KS, Stein SJ, Kurachi M, Petrovic J, Schwartz GW, Mack EA, Uljon S, Wu WW, DeHart AG, McClory SE, Xu L, Gimotty PA, Blacklow SC, Faryabi RB, Wherry EJ, Jordan MS, Pear WS. 2020. Trib1 regulates T cell differentiation during chronic infection by restraining the effector program. Journal of Experimental Medicine 217. doi:10.1084/jem.20190888

Romero-Moya D, Bueno C, Montes R, Navarro-Montero O, Iborra FJ, López LC, Martin M, Menendez P. 2013. Cord blood-derived CD34+ hematopoietic cells with low mitochondrial mass are enriched in hematopoietic repopulating stem cell function. Haematologica. doi:10.3324/haematol.2012.079244

Ron-Harel N, Ghergurovich JM, Notarangelo G, LaFleur MW, Tsubosaka Y, Sharpe AH, Rabinowitz JD, Haigis MC. 2019. T Cell Activation Depends on Extracellular Alanine. Cell Reports 28:3011-3021.e4. doi:10.1016/j.celrep.2019.08.034

Ron-Harel N, Santos D, Ghergurovich JM, Sage PT, Reddy A, Lovitch SB, Dephoure N, Satterstrom FK, Sheffer M, Spinelli JB, Gygi S, Rabinowitz JD, Sharpe AH, Haigis MC. 2016. Mitochondrial Biogenesis and Proteome Remodeling Promote One-Carbon Metabolism for T Cell Activation. Cell Metabolism 24:104–117. doi:10.1016/j.cmet.2016.06.007

Roy DG, Chen J, Mamane V, Ma EH, Muhire BM, Sheldon RD, Shorstova T, Koning R, Johnson RM, Esaulova E, Williams KS, Hayes S, Steadman M, Samborska B, Swain A, Daigneault A, Chubukov V, Roddy TP, Foulkes W, Pospisilik JA, Bourgeois-Daigneault MC, Artyomov MN, Witcher M, Krawczyk CM, Larochelle C, Jones RG. 2020. Methionine Metabolism Shapes T Helper Cell Responses through Regulation of Epigenetic Reprogramming. Cell Metabolism 31:250-266.e9. doi:10.1016/j.cmet.2020.01.006

Saragovi A, Abramovich I, Omar I, Arbib E, Toker O, Eyal G, Berger M. 2020. Systemic hypoxia inhibits T cell response by limiting mitobiogenesis via matrix substrate-level phosphorylation arrest. eLife 9:1–50. doi:10.7554/eLife.56612

Shao Z, Xu P, Xu W, Li L, Liu S, Zhang R, Liu YC, Zhang C, Chen S, Luo C. 2017. Discovery of novel DNA methyltransferase 3A inhibitors via structure-based virtual screening and biological assays. Bioorganic and Medicinal Chemistry Letters 27:342–346. doi:10.1016/j.bmcl.2016.11.023

Shiraki N, Shiraki Y, Tsuyama T, Obata F, Miura M, Nagae G, Aburatani H, Kume K, Endo F, Kume S. 2014. Methionine metabolism regulates maintenance and differentiation of human pluripotent stem cells. Cell Metabolism 19:780–794. doi:10.1016/j.cmet.2014.03.017

Simsek T, Kocabas F, Zheng J, Deberardinis RJ, Mahmoud AI, Olson EN, Schneider JW, Zhang CC, Sadek HA. 2010. The distinct metabolic profile of hematopoietic stem cells reflects their location in a hypoxic niche. Cell Stem Cell. doi:10.1016/j.stem.2010.07.011

Sinclair L V., Howden AJM, Brenes A, Spinelli L, Hukelmann JL, Macintyre AN, Liu X, Thomson S, Taylor PM, Rathmell JC, Locasale JW, Lamond AI, Cantrell DA. 2018. Antigen receptor control of methionine metabolism in T cells. bioRxiv 1–29. doi:10.1101/499095

Sukumar M, Liu J, Mehta GU, Patel SJ, Roychoudhuri R, Crompton JG, Klebanoff CA, Ji Y, Li P, Yu Z, Whitehill GD, Clever D, Eil RL, Palmer DC, Mitra S, Rao M, Keyvanfar K, Schrump DS, Wang E, Marincola FM, Gattinoni L, Leonard WJ, Muranski P, Finkel T, Restifo NP. 2016a. Mitochondrial Membrane Potential Identifies Cells with Enhanced Stemness for Cellular Therapy. Cell Metabolism 23:63–76. doi:10.1016/j.cmet.2015.11.002

Sukumar M, Liu J, Mehta GU, Patel SJ, Roychoudhuri R, Crompton JG, Klebanoff CA, Ji Y, Li P, Yu Z, Whitehill GD, Clever D, Eil RL, Palmer DC, Mitra S, Rao M, Keyvanfar K, Schrump DS, Wang E, Marincola FM, Gattinoni L, Leonard WJ, Muranski P, Finkel T, Restifo NP. 2016b. Mitochondrial Membrane Potential Identifies Cells with Enhanced Stemness for Cellular Therapy. Cell Metabolism. doi:10.1016/j.cmet.2015.11.002

Tannahill GM, Curtis AM, Adamik J, Palsson-McDermott EM, McGettrick AF, Goel G, Frezza C, Bernard NJ, Kelly B, Foley NH, Zheng L, Gardet A, Tong Z, Jany SS, Corr SC, Haneclaus M, Caffery BE, Pierce K, Walmsley S, Beasley FC, Cummins E, Nizet V, Whyte M, Taylor CT, Lin H, Masters SL, Gottlieb E, Kelly VP, Clish C, Auron PE, Xavier RJ, O’Neill LAJ. 2013. Succinate is a danger signal that induces IL-1β via HIF-1α. Nature 496:238–242. doi:10.1038/nature11986.Succinate

Van Den Broek T, Borghans JAM, Van Wijk F. 2018. The full spectrum of human naive T cells. Nature Reviews Immunology 18:363–373. doi:10.1038/s41577-018-0001-y

Vardhana SA, Hwee MA, Berisa M, Wells DK, Yost KE, King B, Smith M, Herrera PS, Chang HY, Satpathy AT, van den Brink MRM, Cross JR, Thompson CB. 2020. Impaired mitochondrial oxidative phosphorylation limits the self-renewal of T cells exposed to persistent antigen. Nature Immunology 21:1022–1033. doi:10.1038/s41590-020-0725-2

Vrisekoop N, den Braber I, de Boer AB, Ruiter AFC, Ackermans MT, van der Crabben SN, Schrijver EHR, Spierenburg G, Sauerwein HP, Hazenberg MD, de Boer RJ, Miedema F, Borghans JAM, Tesselaar K. 2008. Sparse production but preferential incorporation of recently produced naive T cells in the human peripheral pool. Proceedings of the National Academy of Sciences. doi:10.1073/pnas.0709713105

Wang R, Green DR. 2012. Metabolic checkpoints in activated T cells. Nature Immunology 13:907–915. doi:10.1038/ni.2386

Yu G, Wang LG, Han Y, He QY. 2012. ClusterProfiler: An R package for comparing biological themes among gene clusters. OMICS A Journal of Integrative Biology 16:284–287. doi:10.1089/omi.2011.0118

Yu G, Wang LG, He QY. 2015. ChIP seeker: An R/Bioconductor package for ChIP peak annotation, comparison and visualization. Bioinformatics 31:2382–2383. doi:10.1093/bioinformatics/btv145

Yu Q, Zhou B, Zhang Y, Nguyen ET, D. J, Glosson NL, Kaplan MH. 2012. DNA methyltransferase 3a limits the expression of interleukin-13 in T helper 2 cells and allergic airway inflammation. Proceedings of the National Academy of Sciences of the United States of America 109:541–546. doi:10.1073/pnas.1103803109

Yu YR, Imrichova H, Wang H, Chao T, Xiao Z, Gao M, Rincon-Restrepo M, Franco F, Genolet R, Cheng WC, Jandus C, Coukos G, Jiang YF, Locasale JW, Zippelius A, Liu PS, Tang L, Bock C, Vannini N, Ho PC. 2020. Disturbed mitochondrial dynamics in CD8+ TILs reinforce T cell exhaustion. Nature Immunology 21:1540–1551. doi:10.1038/s41590-020-0793-3

Zhang N. 2018. Role of methionine on epigenetic modification of DNA methylation and gene expression in animals. Animal Nutrition 4:11–16. doi:10.1016/j.aninu.2017.08.009

Zhang Y, Liu T, Meyer CA, Eeckhoute J, Johnson DS, Bernstein BE, Nussbaum C, Myers RM, Brown M, Li W, Shirley XS. 2008. Model-based analysis of ChIP-Seq (MACS). Genome Biology 9. doi:10.1186/gb-2008-9-9-r137

